# Single cell, whole embryo phenotyping of pleiotropic disorders of mammalian development

**DOI:** 10.1101/2022.08.03.500325

**Authors:** Xingfan Huang, Jana Henck, Chengxiang Qiu, Varun K. A. Sreenivasan, Saranya Balachandran, Rose Behncke, Wing-Lee Chan, Alexandra Despang, Diane E. Dickel, Natja Haag, Rene Hägerling, Nils Hansmeier, Friederike Hennig, Cooper Marshall, Sudha Rajderkar, Alessa Ringel, Michael Robson, Lauren Saunders, Sanjay R. Srivatsan, Sascha Ulferts, Lars Wittler, Yiwen Zhu, Vera M. Kalscheuer, Daniel Ibrahim, Ingo Kurth, Uwe Kornak, David R. Beier, Axel Visel, Len A. Pennacchio, Cole Trapnell, Junyue Cao, Jay Shendure, Malte Spielmann

## Abstract

Mouse models are a critical tool for studying human diseases, particularly developmental disorders, as well as for advancing our general understanding of mammalian biology. However, it has long been suspected that conventional approaches for phenotyping are insufficiently sensitive to detect subtle defects throughout the developing mouse. Here we set out to establish single cell RNA sequencing (sc-RNA-seq) of the whole embryo as a scalable platform for the systematic molecular and cellular phenotyping of mouse genetic models. We applied combinatorial indexing-based sc-RNA-seq to profile 101 embryos of 26 genotypes at embryonic stage E13.5, altogether profiling gene expression in over 1.6M nuclei. The 26 genotypes include 22 mouse mutants representing a range of anticipated severities, from established multisystem disorders to deletions of individual enhancers, as well as the 4 wildtype backgrounds on which these mutants reside. We developed and applied several analytical frameworks for detecting differences in composition and/or gene expression across 52 cell types or trajectories. Some mutants exhibited changes in dozens of trajectories (*e.g.*, the pleiotropic consequences of altering the *Sox9* regulatory landscape) whereas others showed phenotypes affecting specific subsets of cells. We also identify differences between widely used wildtype strains, compare phenotyping of gain vs. loss of function mutants, and characterise deletions of topological associating domain (TAD) boundaries. Intriguingly, even among these 22 mutants, some changes are shared by heretofore unrelated models, suggesting that developmental pleiotropy might be “decomposable” through further scaling of this approach. Overall, our findings show how single cell profiling of whole embryos can enable the systematic molecular and cellular phenotypic characterization of mouse mutants with unprecedented breadth and resolution.

## Introduction

For over 100 years, the laboratory mouse (*Mus musculus*) has served as the quintessential animal model for studying both common and rare human diseases^1–4^. For developmental disorders in particular, mice have been transformative, as a mammalian system that is nearly ideal for genetic analysis and in which the embryo is readily accessible^5^.

In the first decades of the field, mouse genetics relied on spontaneous or induced mutations resulting in visible physical defects that could then be mapped. However, gene-targeting techniques subsequently paved the way for “reverse genetics”, *i.e.* analysing the phenotypic effects of intentionally engineered mutations. Through systematic efforts such as the International Knockout Mouse Consortium, knockout models are now available for thousands of genes^6^. Furthermore, with the emergence of CRISPR/Cas genome editing^7,8^, it is increasingly practical to delete individual regulatory elements or otherwise modify the *cis-*regulatory landscape, and to then study the *in vivo* consequences of these alterations^9,10^.

Phenotyping has also grown more sophisticated. Conventional investigations of developmental syndromes typically focus on one organ system at a specific stage of development, *e.g.* combining expression analyses, histology, and imaging to investigate a visible malformation^1,11,12^. However, pleiotropy is a pervasive phenomenon in mammalian development, and focusing on one aspect of a phenotype may come at the expense of detecting or characterising others, particularly if they are subtle or masked by lethality. The concept of the Mouse Clinic, in which a given model is subjected to a battery of standardised tests, reflects a more systematic approach^13^. However, such clinics are expensive and time-consuming to conduct in practice. Furthermore, many kinds of phenotypes detected through such tests (*e.g.*, behavioural, electrophysiological) may require years of additional work to link to their molecular and cellular correlates. It is also the case that knockouts of even highly conserved coding or regulatory sequences frequently result in no detectable abnormality or only minor transcriptional changes^14–16^. In such instances, it remains unknown whether there is truly no phenotype, or whether the methods used are simply insufficiently sensitive. In sum, phenotyping has become “rate limiting” in mouse genetics.

The recent emergence of single cell molecular profiling technologies (*e.g.*, sc-RNA-seq) offer a potential path to overcome this barrier. As a first step, we and others have extensively applied sc-RNA-seq to profile wildtype mouse development at the scale of the whole embryo^17–22^. Applying sc-RNA-seq to mouse mutants, several groups have successfully unravelled how specific mutations affect transcriptional networks and lead to altered cell fate decisions in individual organs^23–26^. However, there is still no clear framework for analysing such data at the scale of the whole embryo, nor for how such data from multiple mutants might be combined to better understand the molecular and cellular basis of classic phenomena like pleiotropy.

Here we set out to establish sc-RNA-seq of whole embryos as a scalable framework for the systematic molecular and cellular phenotyping of mouse genetic models. We profiled 101 embryos of 22 different mouse mutants and 4 wildtype backgrounds at E13.5. The resulting mouse mutant cell atlas (MMCA) includes over 1.6M sc-RNA-seq profiles. To analyze these data, we develop and apply new strategies for detecting differences in composition and/or gene expression across 52 cell types or trajectories spanning the whole mid-gestational embryo.

### Single-cell RNA-seq of 101 mouse embryos

We collected a total of 103 mouse embryos, including 22 different mutants and four wildtype (WT) strains (C57BL/6J, G4, FVB, and BALB/C) at embryonic stage E13.5, and generally four replicates per strain (**Fig. 1a**). The mouse mutants were chosen to represent a spectrum of phenotypes ranging from very severe pleiotropic developmental disorders (*e.g.*, *Sox9*, which we expected to affect many organ systems) to knockouts of individual, noncoding regulatory elements (many of which we expected to result in, at best, subtle defects).

**Figure 1.**
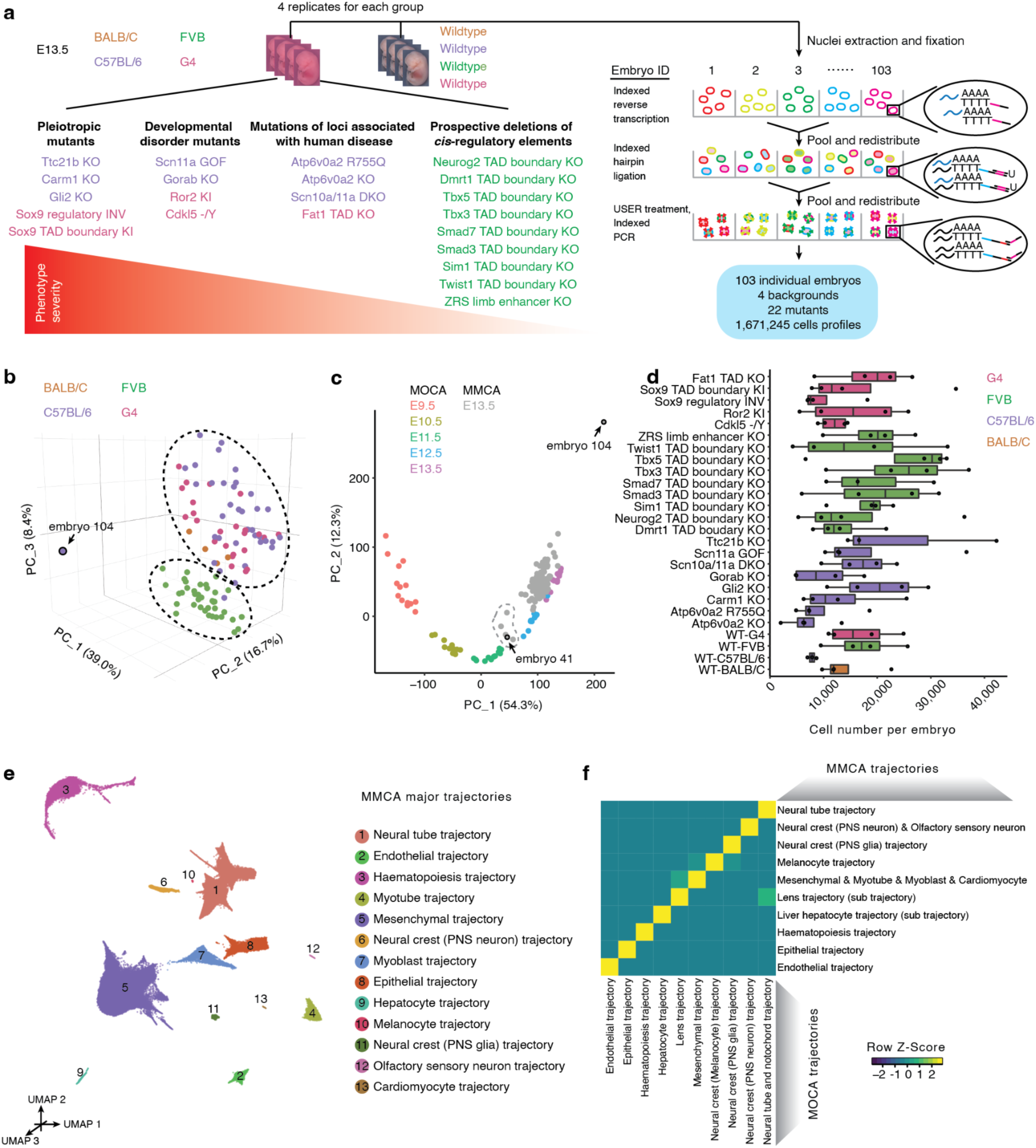
Single-cell RNA-seq of 103 whole mouse embryos staged at E13.5. **a,** We applied sci-RNA-seq3 to profile 1.6M single cell transcriptomes from 103 individual E13.5 embryos, derived from 22 mutants and four wildtype strains, in one experiment. **b,** Embeddings of pseudobulk RNA-seq profiles of MMCA mouse embryos in PCA space with visualisation of top three PCs. Briefly, single cell data from individual embryos of MMCA were aggregated to create 103 pseudobulk samples. Embryos are colored by background strain. The black dotted circles highlight two major groups corresponding to FVB vs. other backgrounds. Embryo #104 was a clear outlier. **c,** Embeddings of pseudobulk RNA-seq profiles of MOCA^17^ and MMCA mouse embryos in PCA space defined solely by MOCA, with MMCA embryos (gray) projected onto it. The top two PCs are visualised. Colored points correspond to MOCA embryos of different stages (E9.5-E13.5), and grey points to MMCA embryos (E13.5). Embryos #104 and #41 were labelled as outliers and removed from the dataset, as discussed in the text. The dashed line (manually added) highlights five MMCA embryos which are colocalized with E11.5 or E12.5 embryos from MOCA. Three are *Scn11a* GOF (#33, #34, #36), one is *Carm1* KO (#101), and one is C57BL/6 wildtype (#41). **d,** The number of cells profiled per embryo for each strain. The centre lines show the medians; the box limits indicate the 25th and 75th percentiles; the replicates are represented by the dots. **e,** 3D UMAP visualisation of wildtype subset of MMCA dataset (215,575 cells from 15 wildtype E13.5 embryos). Cells are colored by major trajectory annotation. **f,** Correlated developmental trajectories between MOCA^17^ and MMCA based on non-negative least-squares (*NNLS*) regression (**Methods**). Shown here is a heat map of the combined regression coefficients (row-scaled) between 10 developmental trajectories from MMCA (rows) and 10 corresponding developmental trajectories from the MOCA (columns). PNS: peripheral nervous system.

We grouped the 22 mutants, all homozygous, into four rough categories (**Supplementary Table 1**): 1) <u>pleiotropic mutants</u>, representing knockouts of developmental genes expressed in multiple organs (*Ttc21b* KO, *Carm1* KO, *Gli2* KO), as well as two mutations of the *Sox9* regulatory landscape suspected to have pleiotropic effects, both of which effectively result in the introduction of a boundary element between endogenous *Sox9* enhancers and the *Sox9* promoter (*Sox9* TAD boundary KI; *Sox9* regulatory INV)^27–30^. 2) <u>developmental disorder mutants</u>, intended to model specific human diseases (*Scn11a* GOF, *Ror2* KI, *Gorab* KO, *Cdkl5 -/*Y)^31–33^,3) <u>mutations of loci</u> <u>associated with human disease</u> (*Scn10a/Scn11a* DKO, *Atp6v0a2* KO, *Atp6v0a2* R755Q, *Fat1TAD* KO)^34,35^. 4) <u>prospective deletions of *cis-*regulatory elements</u>, including of TAD boundaries in the vicinity of developmental transcription factors including *Smad3*, *Twist1*, *Tbx5*, *Neurog2*, *Sim1*, *Smad7*, *Dmrt1*, *Tbx3*, and *Twist1*^36^, and, as a positive control, the ZRS distal enhancer (<u>Z</u>one of polarizing activity <u>R</u>egulatory <u>S</u>equence) which regulates sonic hedgehog (SHH) expression and results in absent distal limb structures^37^.

The 103 flash-frozen embryos (26 genotypes x 4 replicates; one embryo was lost in transport), all staged at E13.5, were sent by five groups to a single site, where they were subjected to sci-RNA-seq3 as previously described^17^. After removing potential doublets, we profiled 1,671,245 nuclei altogether (16,226 +/- 9,289 per embryo; 64,279 +/- 18,530 per strain; median UMI count of 843 per cell and median genes detected of 534 at 75% duplication rate).

Applying principal components analysis (PCA) to “pseudobulk” profiles of the 103 embryos resulted in two roughly clustered groups corresponding to genetic background (**Fig. 1b**). In particular, wildtype and mutant FVB embryos clustered separately from C57BL/6J, G4, and BALB/C embryos. However, embryos corresponding to individual mutants did not cluster separately, suggesting that none were affected with severe, global aberrations and highlighting the inadequacy of bulk RNA-seq for detecting mutant-specific effects. A single outlier embryo (#104) was aberrant with respect to cell recovery (n = 1,047) as well as appearance (**Supplementary Fig. 1**).

We next sought to validate the staging of these embryos, leveraging our previous <u>m</u>ouse <u>o</u>rganogenesis <u>c</u>ell <u>a</u>tlas (MOCA), which spans E9.5 to E13.5^17^. PCA of pseudobulk profiles of 61 wildtype embryos from MOCA resulted in a first component (PC1) that was strongly correlated with developmental age (**Fig. 1c**). Projecting pseudobulk profiles of the 103 MMCA embryos to this embedding resulted in the vast majority of MMCA embryos clustering with E13.5 embryos from MOCA along PC1, consistent with accurate staging. However, five embryos from MMCA appeared closer to E11.5 or E12.5 embryos from MOCA. Four of these were retained as their delay might be explained by their mutant genotype, while one from a wildtype background (C57BL/6; #41) was designated as a second outlier. We removed cells from the two outlier embryos (#104; #41) as well as cells with high proportions of reads mapping to the mitochondrial genome (>10%) or ribosomal genes (>5%). This left 1,627,857 cells, derived from 101 embryos (**Fig. 1d**).

To facilitate an integrated analysis, we sought to project cells from all genotypes to a wildtype derived “reference embedding” (**Supplementary Fig. 2**; **Methods**). We first applied principal components (PC) dimensionality reduction to cells from wildtype genotypes only (n = 215,575; 13.2% of dataset). We then projected cells from mutant genotypes to this embedding, followed by alignment on the combined data to mitigate the effects of technical factors. Next, we applied the UMAP algorithm to the aligned principal components of wildtype cells, followed by Louvain clustering and manual annotation of the resulting major trajectories and sub-trajectories based on marker gene expression. Finally, we projected mutant cells into this UMAP space and assigned them major trajectory and sub-trajectory labels via a *k*-nearest neighbour (*k*-NN) heuristic.

Altogether, we identified 13 major trajectories, 8 of which could be further stratified into 59 sub-trajectories (**Fig. 1e**; **Supplementary Fig. 3**; **Supplementary Table 2**). These were generally consistent with our annotations of MOCA, albeit with some corrections as we have described elsewhere^38,39^, as well as greater granularity for some cell types that is likely a consequence of the deeper sampling of E13.5 cells in these new data (**Fig. 1f**; **Supplementary Fig. 4**). For example, what we had previously annotated as the excitatory neuron trajectory could be further stratified into a di/mesencephalon (*Slc17a6*+, *Barhl1*+, *Shox2*+), thalamus (*Ntng1*+, *Gbx2*+) and spinal cord (*Ebf1*+, *Ebf3*+) sub-trajectories, while skeletal muscle could be further stratified into myoblast (*Pax7*+) and myotube (*Myh3*+, *Myog*+) sub-trajectories.

### Mutant-specific differences in cell type composition

Analogous to how there are many assays for phenotyping a mouse, there are many computational strategies that one might adopt in order to investigate mutant-specific differences in these embryo-scale sc-RNA-seq data. Here we pursued three main approaches: 1) quantification of gross differences in cell type composition (this section); 2) investigation of more subtle differences in the distribution of cell states <u>within</u> annotated trajectories and sub-trajectories; and 3) analysis of the extent to which phenotypic features are shared between mutants.

To systematically assess cell type compositional differences, we first examined the proportions of cells assigned to each of the 13 major trajectories across the 4 wildtype and 22 mutant strains. For the most part, these proportions were consistent across genotypes (**Supplementary Fig. 5a**). However, some mutants exhibited substantial differences. For example, compared to the C57BL/6 wildtype, the proportion of cells falling in the neural tube trajectory decreased from 37.3% to 33.7% and 32.6% in the *Gli2* KO and *Ttc21b* KO mice, respectively, while the proportion of cells falling in the mesenchymal trajectory decreased from 44.1% to 37.1% in the *Gorab* KO mice. These changes are broadly consistent with the gross phenotypes associated with these mutations^28,33,40,41^, but are caveated by substantial interindividual heterogeneity within each genotype (**Supplementary Fig. 5b**). Also of note, we observe differences in major trajectory composition between the four wildtype strains. For example, relative to BALB/C and C57BL/6, the FVB and G4 wildtype mice consistently had substantially lower proportions of cells in the mesenchymal trajectory and higher proportions of cells in the neural tube trajectory (**Supplementary Fig. 5c**).

To increase resolution, we sought to investigate compositional differences at the level of sub- trajectories. For each combination of background (C57BL/6, FVB, G4) and sub-trajectory (n = 54), we performed a regression analysis to identify instances where a particular mutation was nominally predictive of the proportion of cells falling in that sub-trajectory (uncorrected *p-*value < 0.05; beta-binomial regression; **Methods**). Across the 22 mutants, this analysis highlighted 300 nominally significant changes (**Fig. 2a**; **Supplementary Table 3**). Due to the limited number of replicate embryos per wildtype and mutant strain, our power to detect changes is limited, particularly in the smaller trajectories. Nevertheless, several patterns were clear:

**Figure 2.**
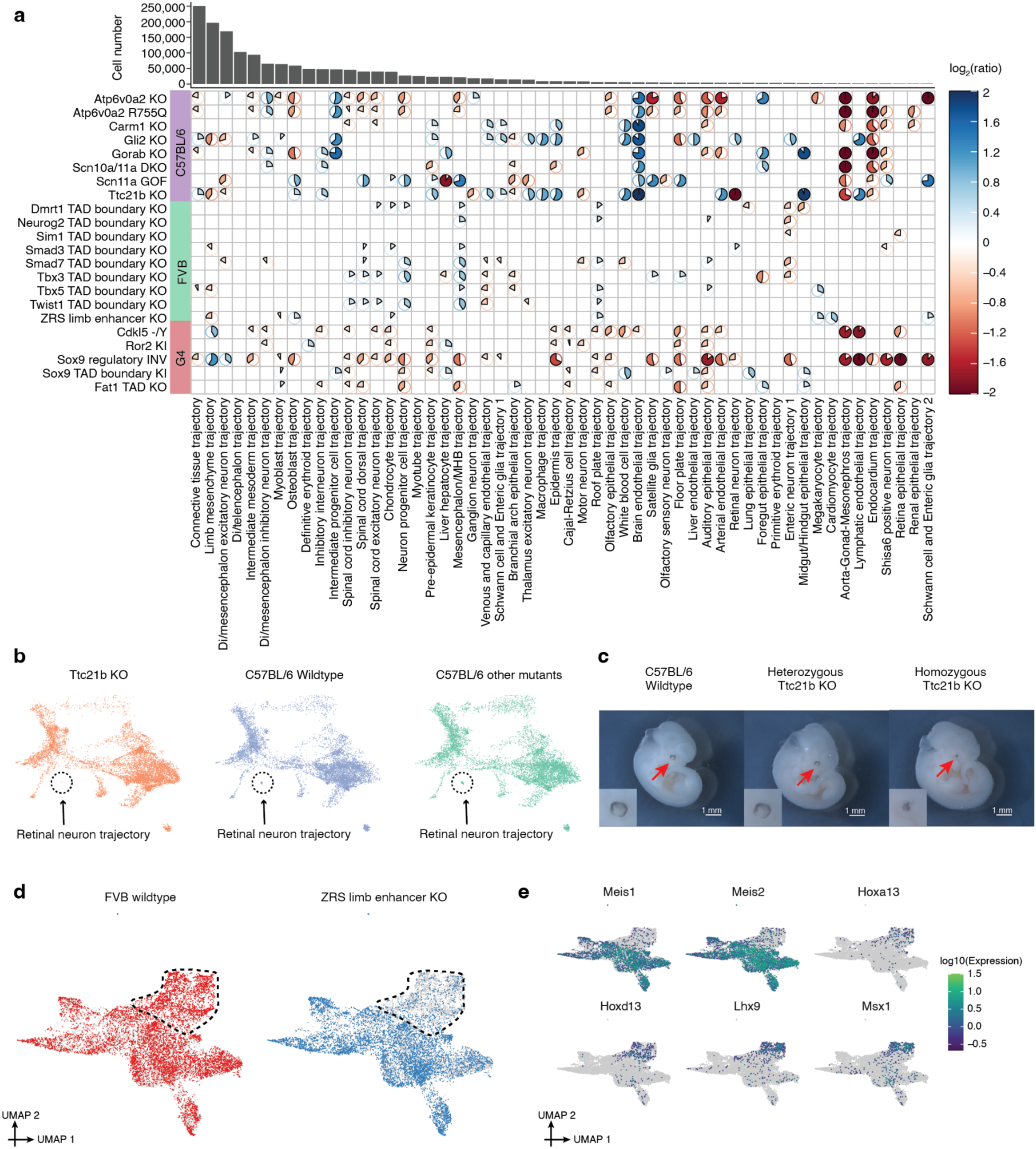
Cell composition changes for individual mutants across developmental trajectories. **a,** Heatmap shows log2 transformed ratios of the cell proportions between each mutant type (*y*-axis) and its corresponding wildtype background, across individual sub-trajectories (*x*-axis). Sub-trajectories with a mean number of cells across individual embryos of less than ten were excluded from this analysis, leaving 54 (columns). Only those combinations of mutant and sub-trajectory which were nominally significant in the regression analysis are shown (see text and **Methods**; uncorrected *p*-value < 0.05; beta-binomial regression). For calculating the displayed ratios, cell counts from replicates were merged. The pie color and direction correspond to whether the log2 transformed ratio is above 0 (blue, clockwise) or below 0 (red, anticlockwise), while the pie size and colour intensity correspond to the scale of log2 transformed ratio. A handful of log2 transformed ratios with > 2 (or < -2) were manually set to 2 (or -2) for a better visualisation. The number of cells assigned to each developmental trajectory in the overall dataset is shown above the heatmap. **b,** 3D UMAP visualisation of the neural tube trajectory, highlighting cells from either the *Ttc21b* KO (left), C57BL/6 wildtype (middle), or other mutants on the C57BL/6 background (right). The three plots were randomly downsampled to the same number of cells (*n* = 8,749 cells). **c,** Homozygous *Ttc21b* KO mice embryo (E11.5) showed abnormal eye development. **d,** UMAP visualisation of co-embedded cells of limb mesenchyme trajectory from the ZRS limb enhancer KO and FVB wildtype. The same UMAP is shown twice for both, highlighting cells from either FVB wildtype (left) or ZRS limb enhancer KO (right). The subset of cells in this co-embedding exhibiting more extreme loss in the ZRS limb enhancer KO is highlighted. **e,** The same UMAP as in panel **d**, colored by gene expression of marker genes which appear specific to proximal limb development (*Meis1*, *Meis2*)^48,49^ and distal limb development (*Hoxa13, Hoxd13, Lhx9, Msx1*)^47,50,51^. Gene expression was calculated from original UMI counts normalised to size factor per cell, followed by 10-log transformation. PNS: peripheral nervous system. MHB: midbrain-hindbrain boundary. Di: Diencephalon.

First, it is evident that *Atp6v0a2* KO and *Atp6v0a2* R755Q, two distinct mutants of the same gene^34^, are assigned very similar patterns by this analysis, both with respect to which sub-trajectories are nominally significant as well as the direction and magnitude of changes (first two rows of **Fig. 2a**). Although perhaps expected, the consistency supports the validity of this analytical approach.

Second, the mutants varied considerably with respect to the number of sub-trajectories that were nominally significant for compositional differences. At the higher extreme, the proportions of cells falling in 30 of 54 sub-trajectories were nominally altered by the *Sox9* regulatory INV mutation, consistent with the wide-ranging roles of Sox9 in development^42,43^. On the other hand, other mutants, such as the TAD boundary knockouts, exhibited comparatively few changes, consistent with the paucity of gross phenotypes in such mutants^16^. Nonetheless, all TAD boundary knockouts did show some changes, including specific ones, *e.g.* the lung epithelial and liver hepatocyte trajectories were decreased in the *Dmrt1* and *Tbx3* TAD boundary KOs, respectively, but not in other TAD boundary knockouts. At the lower extreme, the *Sim1* TAD boundary KO exhibited just two altered sub-trajectories.

Third, some sub-trajectories exhibited altered proportions in many mutants (*e.g.* the mesencephalon/MHB trajectory in 12 mutants) while others were changed only in a few (*e.g.* the definitive erythroid trajectory in *Ror2* KI only). In some cases, such patterns were “block-like” by background strain (*e.g.* all B6 mutants exhibited gains in endothelial cells and losses in endocardium). Although particular sub-trajectories might be vulnerable to disruption in a strain- specific way, it is also possible that this is a technical artefact (*e.g.* if the four wildtype replicates that we profiled for a given strain were atypical).

There were a few extreme examples, *e.g.* where a sub-trajectory appeared to be fully lost in a specific mutant. For example, *Ttc21b*, which encodes a cilial protein and whose knockout is associated with brain, bone and eye phenotypes^28,44,45^, exhibited a dramatic reduction in the proportion of cells in the retinal neuron trajectory (log_2_(ratio) = -6.69; unadjusted *p*-value = 0.028; beta-binomial regression) (**Fig. 2b**), as well as the lens (log_2_(ratio) = -2.64) and retina epithelium (log_2_(ratio) = -2.32) trajectories (Supplementary Fig. 6). Validating this finding, the developing eye appears diminished in the homozygous *Ttc21b* mutant at E11.5 embryos compared to the wildtype or heterozygous mutant (**Fig. 2c**).

However, most changes were relatively subtle. For example, the ZRS limb enhancer KO is a well- studied mutant which shows a loss of the distal limb structure at birth^46^. This analytical framework highlighted eight sub-trajectories whose proportions were nominally altered in the ZRS limb enhancer KO, most of which were mesenchymal. However, although the most extreme, the reduction in limb mesenchymal cells was only about 30% (log_2_(ratio) = -0.49; unadjusted *p*-value = 6.32e-3; beta-binomial regression). To assess whether further subpopulations of the limb mesenchyme were more substantially changed, we performed co-embedding of limb mesenchyme cells from the ZRS limb enhancer KO and the FVB wildtype. Indeed, a subpopulation of the limb mesenchyme was much more markedly affected (**Fig. 2d**; **Supplementary Fig. 7a**), and this subpopulation specifically expressed markers of the distal mesenchyme of the early embryonic limb bud, such as *Hoxa13* and *Hoxd13* (**Fig. 2e**)^47^. Of note, we did not observe such heterogeneity when we examined the seven other sub-trajectories whose proportions were nominally altered in the ZRS limb enhancer KO (**Supplementary Fig. 7b**), consistent with the specificity of this phenotype.

### LochNESS analysis reveals differences in transcriptional state within cell type trajectories

Given that most of the mutants that we studied did not exhibit macroscopic anatomical defects or otherwise severe phenotypes at E13.5, we next sought to develop a more sensitive approach for detecting deviations in transcriptional programs <u>within</u> cell type trajectories. Specifically, we developed “lochNESS” (<u>lo</u>cal <u>c</u>ellular <u>h</u>euristic <u>N</u>eighbourhood <u>E</u>nrichment <u>S</u>pecificity <u>S</u>core), score that is calculated based on the “neighbourhood” of each cell in a sub-trajectory co- embedding of a given mutant (all replicates) vs. a pooled wildtype (all replicates of all backgrounds) (**Fig. 3a; Methods**; although developed independently, this approach is similar to recent work by Dann and colleagues^52^). Briefly, we took the aligned PC features of each sub-trajectory, as described above, and found k-NNs for each cell, excluding cells from the same mutant replicate from consideration. For each mutant cell, we then computed the ratio of the observed vs. expected number of mutant cells in its neighbourhood, with expectation simply based on the overall ratio of mutant vs. wildtype cells in co-embedding. In the scenario where mutant and wildtype cells are fully mixed, the resulting ratio should be close to 1. The final lochNESS was defined as the ratio minus 1, equivalent to the fold change of mutant cell composition.

**Figure 3.**
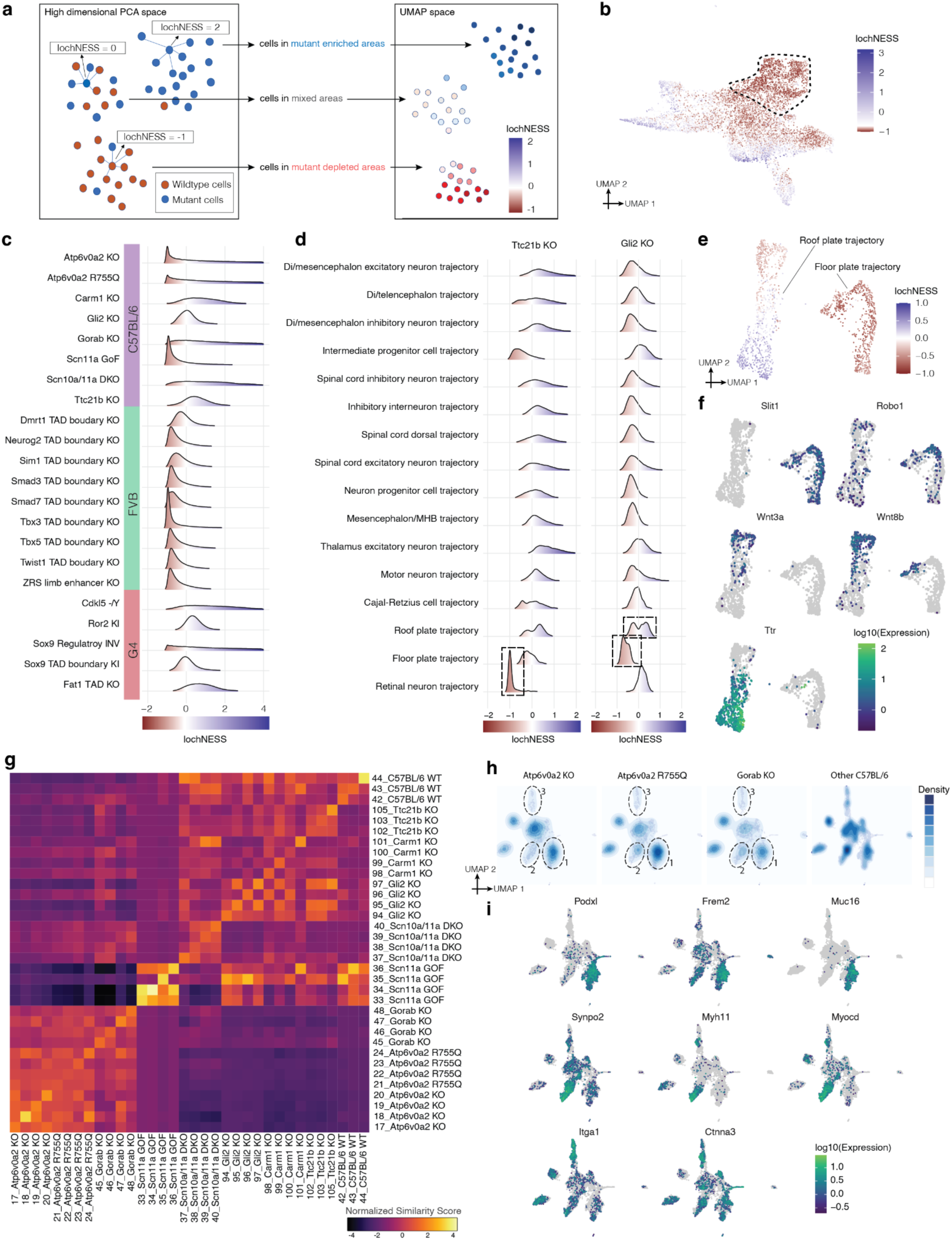
LochNESS analysis identifies mutant related changes. **a,** Schematic of lochNESS calculation and visualisation. **b,** UMAP visualisation of co-embedded cells of limb mesenchyme trajectory from the ZRS limb enhancer KO and FVB wildtype, colored by lochNESS, with colour scale centred at the median of lochNESS. The subset of cells in this co-embedding that corresponds to the area exhibiting more extreme loss in the ZRS limb enhancer KO cells in Fig. 2d is highlighted (dashed circle). **c,** Distribution of lochNESS across all 64 sub-trajectories in each mutant. **d,** Distribution of lochNESS in the neural tube sub-trajectories of the *Ttc21b* KO and *Gli2* KO mutants. Dashed boxes highlight the shifted distributions of the retinal neuron sub-trajectory of the *Ttc21b* KO mutant and the floor plate and roof plate sub-trajectories of the *Gli2* KO mutant. **e,** UMAP visualisation of co-embedded cells of the floor plate and roof plate sub-trajectories from the Gli2 KO mutant and pooled wildtype, colored by lochNESS. **f,** same as in panel **e**, but colored by expression of selected marker genes. **g,** Heatmap showing similarity scores between individual C57BL/6 embryos in the mesenchymal trajectory. Rows and columns are grouped by genotype and labelled by embryo id and genotype. **h,** UMAPs showing the co-embedding of the intermediate mesoderm sub- trajectory for mutants from the C57BL/6 background strain, with cell density and distributions overlaid. Dashed circles highlight three clusters of cells where *Atp6v0a2* KO, *Atp6v0a2* R755Q and *Gorab* KO mice exhibit enrichment (cluster 1) or depletion (clusters 2 & 3), compared to other mutants in the C57BL/6 background strain. **i,** same as in panel **h**, but colored by expression of marker genes of the clusters highlighted in panel **h**.

Visualisation of lochNESS in the embedded space highlights areas with enrichment or depletion of mutant cells. For example, returning to the previously discussed ZRS limb enhancer KO mice, we observed markedly low lochNESS in a portion of the limb mesenchymal trajectory corresponding to the distal limb (**Fig. 3b**; **Fig. 2d**). This highlights the value of the lochNESS framework, as within the sub-trajectory (limb mesenchyme), an effect could be detected and also assigned to a subset of cells in a label-agnostic fashion.

Plotting the global distributions of lochNESS for each mutant across all sub-trajectories, we further observed that some mutants (*e.g.* most TAD boundary knockouts; *Scn11a* GOF) exhibit unremarkable distributions (**Fig. 3c**). However, others (*e.g. Sox9* regulatory INV; *Scn10a/11a* DKO) are associated with a marked excess of high lochNESS, consistent with mutant-specific effects on transcriptional state across many developmental systems. Of note, we confirmed that repeating the calculation of lochNESS after random permutation of mutant and wildtype labels resulted in bell-shaped distributions centred around zero (**Supplementary Fig. 8a**). As such, the deviance of lochNESS can be summarised as the average euclidean distance between lochNESS vs lochNESS under permutation (**Supplementary Fig. 8b**).

We next examined lochNESS within each mutant of each sub-trajectory to identify system-specific phenotypes. For example, consistent with results shown above, we observed low lochNESS within the retinal neuron sub-trajectory in the *Ttc21b* KO (**Fig. 3d**; **Supplementary Fig. 8c**). We also observed a strong shift towards low scores for the floor plate sub-trajectory in the *Gli2* KO, and interestingly, a more subtle change in lochNESS distribution for the roof plate trajectory, which is forming opposite to the floor plate along the D-V axis of the developing neural tube (**Fig. 3d**; **Supplementary Fig. 8c**). To explore this further, we extracted and reanalyzed cells corresponding to the floor plate and roof plate. Within the floor plate, *Gli2* KO cells consistently exhibited low lochNESS (**Fig. 3e**). However, there were only a handful of differentially expressed genes between wildtype and mutant cells, and no significantly enriched pathways within that set. For example, genes like *Robo1* and *Slit1*, both involved in neuronal axon guidance, are specifically expressed in the floor plate relative to the roof plate (**Fig. 3f**; **Supplementary Fig. 8g**), but are not differentially expressed between wildtype and *Gli2* KO cells of the floor plate. Alternatively, our failure to detect substantial differential expression may be due to power, as there were fewer floor plate cells in the *Gli2* KO (∼60% reduction). Overall, these observations are consistent with the established role of Gli2 in floor plate induction and the previous demonstration that *Gli2* knockouts fail to induce a floor plate (Matise et al. 1998; Ding et al. 1998).

Less expectedly, this focused analysis also revealed two subpopulations of roof plate cells, one depleted and the other enriched for *Gli2* KO cells (**Fig. 3e**; **Supplementary Fig. 8d-f**). To annotate these subpopulations, we examined genes whose expression was predictive of lochNESS via regression (**Methods**). The mutant-enriched group of roof plate cells was marked by *Ttr*, a marker for choroid plexus and dorsal roof plate development^53^, as well as genes associated with the development of cilia (*e.g. Cdc20b*, *Gmnc*, *Dnah6* and *Cfap43*), while the mutant-depleted group was marked by Wnt signaling-related genes including *Rspo1/2/3* and *Wnt3a/8b/9a* (**Fig. 3f**; **Supplementary Fig. 8g**; **Supplementary Table 3**)^54–57^. It has been shown that ventrally-expressed Gli2 plays a central role in dorsal-ventral patterning of the neural tube by antagonising Wnt/Bmp signalling from the dorsally-located roof plate^58^. Our results are consistent with this, and also define two subpopulations of roof plate cells on which Gli2 KO appears to have differential effects. Of note, the relatively subtle and opposing effects on these roof plate subpopulations were missed by our original analysis of cell type proportions, and only uncovered by the granularity of the lochNESS strategy.

LochNESS distributions can be systematically screened to identify sub-trajectories exhibiting substantial mutant-specific shifts. For example, while all TAD boundary KO mutants have similarly unremarkable global lochNESS distributions, when we plot these distributions by sub-trajectory, a handful of shifted distributions are evident (**Supplementary Fig. 9a**). Such deviations, summarised as the average euclidean distances between lochNESS and lochNESS under permutation, are visualised in **Supplementary Fig. 9b**. For example, multiple epithelial sub-trajectories, including pre−epidermal keratinocyte, epidermis, branchial arch, and lung epithelial trajectories, are most shifted in *Tbx3* TAD boundary KO cells. Co-embeddings of mutant and wildtype cells of these sub-trajectories, together with regression analysis, identify multiple keratin genes as positively correlated with lochNESS, consistent with a role for *Tbx3* in epidermal development (**Supplementary Fig. 9c-d**; **Supplementary Table 4**)^59,60^. The lung epithelial cells were separated into two clusters, with the cluster more depleted in *Tbx3* TAD boundary KO cells marked by *Etv5*, a transcription factor associated with alveolar type II cell development, as well as *Bmp* signalling genes that regulate *Tbx3* during lung development (*Bmp1/4*), and distal airway markers *Sox9* and *Id2*^60–62^.

### Identification of mutant-specific and mutant-shared effects

Pleiotropy, wherein a single gene influences multiple, unrelated traits, is a pervasive phenomenon in developmental genetics, and yet remains poorly understood^63^. A corollary of pleiotropy is that there are also specific traits that appear to be influenced by multiple, unrelated genes. For the most part, the characterization of the sharing of phenotypic features between multiple Mendelian disorders has remained coarse. For example, many disorders share macrocephaly as a feature, but it remains largely unexplored whether the molecular and cellular basis for macrocephaly is shared between them, unique to each, or somewhere in between.

Although here we have “whole embryo” molecular profiling of just 22 mutants, we sought to investigate whether we could distinguish between mutant-specific and mutant-shared effects within each major trajectory. In brief, within a co-embedding of cells from all embryos from a given background strain, we computed k-NNs as in **Fig. 3a**, and then calculated the observed vs. expected ratio of each genotype among a cell’s k-NNs. The “similarity score” between one genotype vs. all others is defined as the mean of these ratios across cells of the genotype. To assess whether any observed similarities or dissimilarities are robust, we can also calculate similarity scores between individual embryos. For example, for the mesenchymal trajectory of C57BL/6 mutants, similarity scores are generally higher for pairwise comparisons of individuals with the same genotype (**Fig. 3g**; **Supplementary Fig. 10a-b**).

The *Scn11a* GOF mutant exhibited the most extreme similarity scores, in terms of both similarity between replicates and dissimilarity with other genotypes (**Fig. 3g**; **Supplementary Fig. 10a**). The *Scn11a* GOF mutant carries a missense mutation in the *Scn11a* locus which is reported to result in reduced pain sensitivity both in mice and men without obvious signs of neurodegeneration, suggesting altered electrical activity of peripheral pain-sensing neurons and impaired synaptic transmission to postsynaptic neurons (Leipold et al. 2013). However, at least grossly, the mutant does not seem to be associated with mesenchymal phenotypes. Noting that the *Scn11a* GOF mutant embryos clustered with E12.5 embryos instead of E13.5 embryos in our pseudobulk analysis (**Fig. 1c**), we speculated that its extreme similarity scores might be attributable to developmental delay of the *Scn11a* GOF mutant at the scale of the whole embryo. To investigate this further, we co-embedded *Scn11a* GOF mutant cells with pooled wildtype cells and MOCA cells from the neural tube trajectory. While wildtype cells were distributed near E13.5 cells from MOCA, the *Scn11a* GOF cells were embedded closer to cells from earlier developmental timepoints (**Supplementary Fig. 10d**). As a more systematic approach, we calculated a “time score” for each cell from the MMCA dataset by taking the k-NNs of each MMCA cell in the MOCA dataset and calculating the average of the developmental time of the MOCA cells. The relative time score distributions of *Scn11a* GOF cells and wildtype cells suggest that *Scn11a* GOF cells are significantly delayed in all major trajectories examined (single sided student’s t-test, raw p-value < 0.01; **Supplementary Fig. 10e**). As such, the apparently unique signature of *Scn11a* GOF cells might be attributable to these embryos simply being earlier in development, suggesting a more global role for sodium ion channels not only for neuronal function but also early development and cell fate determination^64^. Incorrect staging is formally possible, but unlikely because the embryos derived from three independent litters.

In sharp contrast with the relative uniqueness of the *Scn11a* GOF mutant, we also observed that the similarity scores between three mutants -- *Atp6v0a2* KO, *Atp6v0a2* R755Q and *Gorab* KO -- was consistent with <u>shared</u> effects, in the mesenchymal, epithelial, endothelial, hepatocyte and neural crest (PNS glia) trajectories in particular; in other main trajectories, such as neural tube and hematopoiesis, *Atp6v0a2* KO and *Atp6v0a2* R755Q exhibited high similarity scores with one another, but not with *Gorab* KO (**Fig. 3g**; **Supplementary Fig. 10a,c,f**). Such sharing is perhaps expected between the *Atp6v0a2* KO and *Atp6v0a2* R755Q mutants, as they involve the same gene. In human patients, mutations in *ATP6V0A2* and *GORAB* cause overlapping connective tissue disorders, which is reflected in the misregulation of the mesenchymal trajectory of *Atp6v0a2* and *Gorab* mutants^34–33^. However, only the ATP6V0A2-related disorder displays a prominent CNS phenotype, consistent with the changes in the neural tube trajectory seen only in both *Atp6v0a2* models (**Supplementary Fig. 10a,c,f**).

In order to explore phenotypic sharing between these genotypically distinct mutants at greater granularity, we co-embedded cells of the intermediate mesoderm sub-trajectory from C57BL/6 strains. We identified three subclusters of intermediate mesoderm where *Atp6v0a2* KO, *Atp6v0a2* R755Q and *Gorab* KO mice are similarly distributed compared to other C57BL/6 genotypes (**Fig. 3h,i**). In particular, cluster 1 is enriched for cells from *Atp6v0a2* KO, *Atp6v0a2* R755Q and *Gorab* KO mice and is marked by genes related to epithelial-to-mesenchymal transition, cell-cell adhesion and migration, such as *Podxl*, *Frem2* and *Muc16* ^65–67^. Clusters 2 and 3 are depleted in cells from *Atp6v0a2* KO, *Atp6v0a2* R755Q and *Gorab* KO mice and are marked by muscular development related genes like *Synpo2*, *Myh11* and *Myocd* (cluster 2), and cell-cell adhesion related genes like *Itga1* and *Ctnna3* (cluster 3)^68–72^.

Altogether, these analyses illustrate how the joint analysis of mutants subjected to whole embryo sc-RNA-seq has the potential to reveal sharing of molecular and cellular phenotypes. This includes global similarity (*e.g. Atp6v0a2* KO vs. *Atp6v0a2* R755Q) as well as instances in which specific aspects of phenotypes are shared between previously unrelated mutants (*e.g. Atp6v0a2* mutants vs. *Gorab* KO).

### Global developmental defects in *Sox9* regulatory mutant

About half of the mutants profiled in this study model disruptions of regulatory, rather than coding, sequences. Among these, the *Sox9* regulatory INV mutant stands out in having a dramatically shifted lochNESS distribution, particularly in the mesenchymal trajectory (**Fig. 3c**; **Fig. 4a**). The *Sox9* locus encodes a pleiotropic transcription factor that plays a central role during the development of the skeleton, the brain, in sex determination as well as several other tissues during embryogenesis, orchestrated by a complex regulatory landscape^73–81^. This particular mutant features an inversion of a 1Mb region upstream of *Sox9* that includes several distal enhancers and a TAD boundary, essentially relocating these elements into a TAD with *Kcnj2*, which encodes a potassium channel (**Fig. 4b**)^27 82,83^ .Consistent with the heterozygous and homozygous *Sox9* knockout, the homozygous *Sox9* regulatory INV is perinatally lethal, with extensive skeletal phenotypes including digit malformation, a cleft palate, bowing of bones and delayed ossification. In addition to the loss of 50% of Sox9 expression, the inversion was previously shown to lead to pronounced missexpression of *Kcnj2* in the digit anlagen in a wildtype *Sox9* pattern^27^. However, the extent to which *Kcnj2* and *Sox9* are mis-expressed elsewhere, as well as the molecular and cellular correlates of the widespread skeletal phenotype, have yet to be deeply investigated.

**Figure 4.**
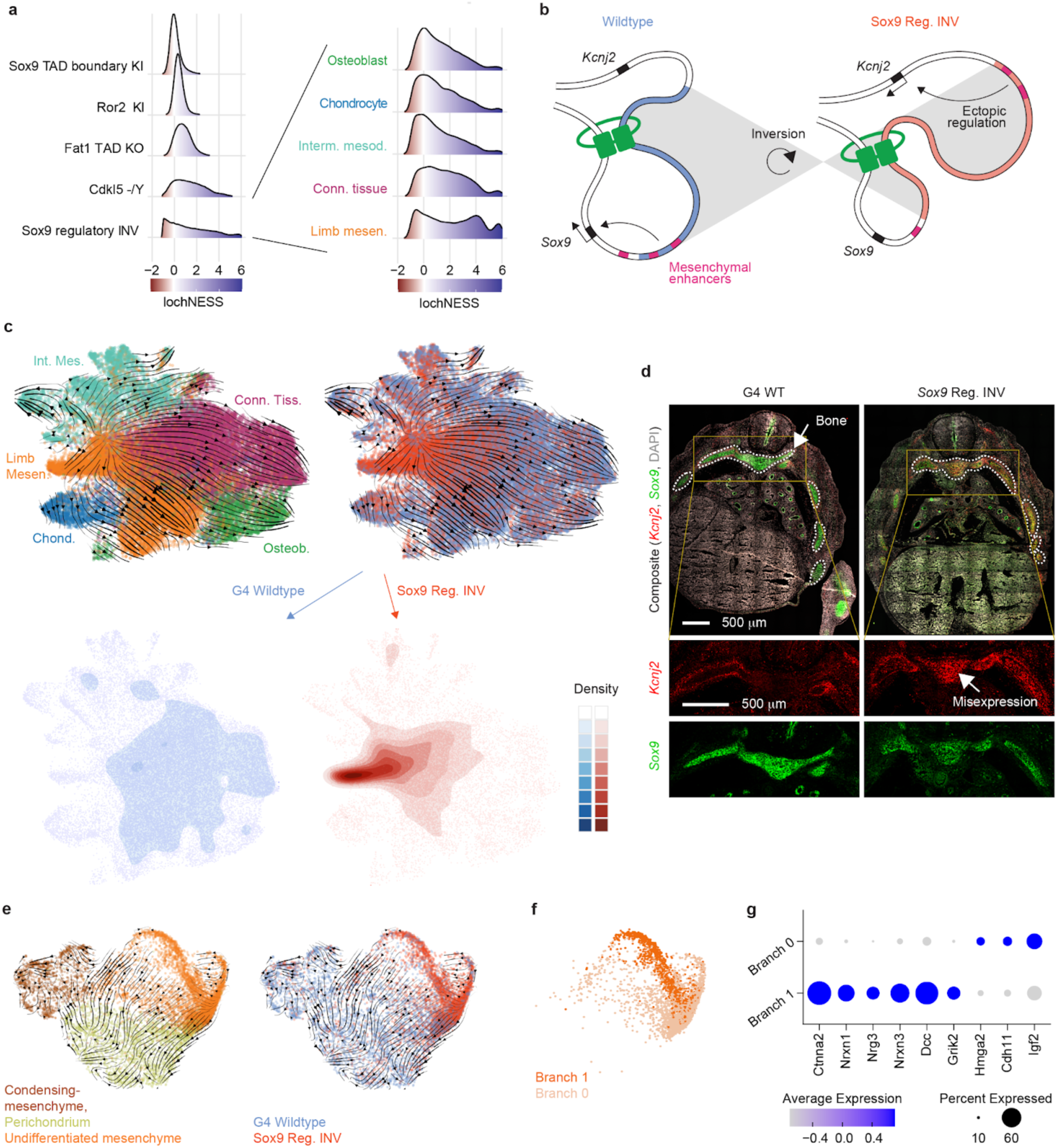
Apparent stalling and redirection of mesenchyme differentiation in the Sox9 regulatory INV mutant. **a,** LochNESS distributions for all G4 mutants in the mesenchymal trajectory (left) and the *Sox9* regulatory INV mutant in mesenchymal sub-trajectories (right). **b,** Model of *Sox9* regulatory INV mutation depicting ectopic *Kcnj2* expression due to adoption of chondrogenesis and osteogenesis specific enhancers. **c,** top: RNA velocity on UMAP embedding of mesenchymal G4 wildtype and *Sox9* regulatory INV cells labelled by annotation (left) or sample (right). bottom: 2D density plots of the same UMAP embedding for G4 wildtype (left) and *Sox9* regulatory INV cells (right). **d,** S*ox9* regulatory INV heterozygous mutant and littermate wildtype RNA scope images (red: *Kcnj2*; green: *Sox9*), with insets below highlighting a region corresponding to developing bone (white circled area) **e**, RNA velocity on UMAP embedding of G4 wildtype and *Sox9* regulatory INV cells in the limb mesenchymal trajectory labelled by annotation (left) or sample (right). **f.** UMAP embedding of *Sox9* regulatory INV cells in the undifferentiated mesenchyme, visualised in the same embedding as in panel **e**. **g**, Dot plot of the top six (where available) significantly differentially expressed genes between the two branches.

At the level of mesenchymal sub-trajectories, shifts in lochNESS distribution for *Sox9* regulatory INV were consistently observed, but the limb mesenchyme and connective tissue were particularly enriched for cells with extremely high lochNESS (**Fig. 4a**, right). Of relevance, 2 of the 3 major enhancers (E250 and E195) known to drive *Sox9*-mediated chondrogenesis in mesenchymal stem cells are located within the inverted region (**Fig. 4b**)^75^. Cell type composition analysis (**Fig. 2a**) showed that *Sox9* regulatory INV mutants harbor considerably larger numbers of cells classified as limb mesenchyme, at the expense of osteoblasts, intermediate mesoderm, chondrocytes and connective tissue trajectory. This shift can also be seen in a UMAP embedding (**Fig. 4c**), a topic that we revisit further below.

These changes in cell type composition were accompanied by reduced expression of *Sox9* and increased expression of *Kcnj2* in bone (aggregate of chondrocyte, osteoblast, limb mesenchyme; **Supplementary Fig. 11a**), although the number of cells expressing *Kcnj2* was generally low. This suggests that the *Sox9* regulatory inversion is resulting in increased *Kcnj2* expression (via *Sox9* enhancer adoption) and *Sox9* reduction (via boundary repositioning) not only in the digit anlagen, but in skeletal mesenchyme more generally. To validate this, we performed RNA *in situ* hybridization (RNAscope) on sections of developing bones of the rib cage at E13.5, comparing a heterozygous *Sox9* regulatory INV mouse with a wildtype littermate. Consistent with our sc-RNA- seq data derived from homozygous mutants, we observe a *Sox9*-patterned increase in *Kcnj2* levels, together with losses in *Sox9* expression, in the developing bone (**Fig. 4d**; **Supplementary Fig. 11b**).

Since the inverted *Sox9* regulatory region also hosts multiple enhancers active in other tissues (*e.g.* E161-lung; E239-cerebral cortex)^75^, we wondered whether these patterns were also seen in other tissues. Indeed, both sc-RNA-seq expression analysis and RNAscope quantification show increased *Kcnj2* levels in all other tissues examined. While reductions in *Sox9* expression, clear in bone, were not observed in most other tissues in our single cell data, RNAscope quantification showed reductions in *Sox9* expression in the telencephalon and lung as well (**Supplementary Fig. 11**). Taken together, these data suggest marked changes in mesenchyme due to reductions of *Sox9* expression (presumably due to separation from key enhancers), together with broader increases in *Kcnj2* expression (presumably due to the appropriation of *Sox9* enhancers).

To explore the apparent effects of the *Sox9* regulatory inversion on mesenchyme in more detail, in particular the apparent accumulation of limb mesenchyme (**Fig. 4c**), we reanalyzed mutant and wildtype cells from the limb mesenchyme sub-trajectory on their own, which revealed subsets corresponding to condensing mesenchyme, perichondrium, and undifferentiated mesenchyme (**Supplementary Fig. 12a,b**). This analysis further revealed that the vast majority of limb mesenchyme “accumulation” in mutant embryos was due to a large proportion of cells that appear delayed or stalled in an undifferentiated or stem-like state, rather than an accumulation of more advanced limb mesenchyme (**Fig. 4c**, bottom panels; **Supplementary Fig. 12a**). Of note, because the annotation of “limb mesenchyme” for this sub-trajectory was propagated forward from earlier stages of development during the creation of MOCA, we cannot rule out that other, non-limb mesenchymal populations contribute to this expanded, undifferentiated pool in the *Sox9* regulatory INV embryos as well.

Inspection of density plots and RNA velocity suggested that wildtype undifferentiated mesenchymal cells (a subset of cells annotated as limb mesenchyme in **Fig. 4c**) are poised to undergo differentiation into diverse subtypes (**Fig. 4c**; **Supplementary Fig. 12a**). In sharp contrast, undifferentiated mesenchymal cells from *Sox9* regulatory INV embryos accumulate at the “source” of differentiation, and also appear to acquire a distinct state (high density region in bottom right sub-panel of **Fig. 4c**). This accumulation is even more apparent in integrated views of the limb mesenchyme sub-trajectory, where we observe two distinct branches, each heavily enriched for *Sox9* regulatory INV mutant cells, within undifferentiated mesenchyme (**Fig. 4e**; **Supplementary Fig. 13a**).

To investigate these two branches further, we performed sub-clustering of *Sox9* regulatory INV undifferentiated mesenchyme cells, followed by differential expression analysis (**Fig. 4f,g**). Interestingly, the most differentially expressed genes in “branch 1” were neuronal, *e.g.* several neurexins and neuregulin 3, an observation that was supported by single-sample gene set enrichment analysis (ssGSEA)^84^, which further highlighted KRAS and other signalling pathways (**Fig. 4g**; **Supplementary Fig. 13b,c**). Of note, mesenchymal stem cells can be differentiated to neuronal states *in vitro*^85^*. A*lthough further investigation is necessary, we note that cells contributing to “branch 0” as well as the neuronal-trending “branch 1” are present in wildtype embryos, albeit at much reduced frequencies compared to the *Sox9* regulatory INV mutant (**Supplementary Fig. 13a**, left).

In sum, consistent with what is known about the role of *Sox9* as a driver gene in cartilage and skeletal development, our data reveals a redirection in the differentiation of osteoblast, chondrocytes and other derivatives of the undifferentiated mesenchyme in the *Sox9* regulatory INV mutant. Among mutants on the G4 background, the observed pattern is specific to the *Sox9* regulatory INV mutant (**Supplementary Fig. 14**). Remarkably however, when we examine the entire dataset (**Supplementary Fig. 14-16**), we observe a similar accumulation of undifferentiated mesenchymal cells in the *Atp6v0a2* KO, *Atp6v0a2* R755Q, and *Gorab* KO mutants, indicating sharing of this sub-phenotype amongst 4 of 22 mutants examined (**Supplementary Fig. 16-17**). This observation further illustrates the potential for systematic, whole embryo analysis to reveal sharing of molecular and cellular sub-phenotypes across pleiotropic developmental mutants in unexpected ways.

## Discussion

In this study, we set out to establish whole embryo sc-RNA-seq as a new paradigm for the systematic, scalable phenotyping of mouse developmental mutants. In one experiment, we generated ∼1.6M single cell transcriptomes from just over 100 E13.5 embryos corresponding to 22 mutant genotypes and 4 wildtype strains. To investigate the resulting dataset, we developed analytical approaches to identify deviations in cell type composition, subtle differences in gene expression within cell types (“lochNESS”), and sharing of sub-phenotypes between mutants (“similarity scores”). We also evaluated how a range of gross phenotypic severities manifest at the molecular and cellular levels, and show how global analysis can in some cases reveal molecular and cellular phenotypes that may be missed by conventional phenotyping. Such “*in silico* developmental biology”, wherein global profiles of developmental mutants are subjected to systematic, outcome-agnostic computational analyses, may complement and guide conventional phenotyping, which can be impractical to scale to all physiological systems even for a single mutant.

We emphasise that the concurrent analysis of many mutants proved essential to the contextualization of particular observations, *i.e.* to understand how specific or non-specific any apparent deviation really was, against a background of dozens of genotypes and over 100 embryos. This aspect of the study also enabled us to discover shared aspects of phenotypes between previously unrelated genotypes, *e.g.* between *Gorab* and *Atp6v0a2* mutants. Looking forward, profiling of additional mouse mutants might enable the further “decomposition” of developmental pleiotropy, a poorly understood phenomenon, into “basis vectors” (*e.g.* the stalling of undifferentiated mesenchyme in 4 of 22 mutants examined).

Our mouse mutant cell atlas (MMCA) has limitations. First, we only profiled 4 replicates per mutant at a single developmental time point. We can’t exclude that some subtle effects were missed that might have been captured through profiling of a larger number of replicate embryos. Second, we profiled only ∼15,000 cells per embryo, which is only a small fraction of the millions of cells that are present in E13.5 embryos, which may also have limited sensitivity. A counterweight to these limitations is that for any given mutant, we had over 1.5M cells from other genotypes (wildtype or other mutants), which facilitated the detection of mutant-specific phenotypes for even rare cell types, *e.g.* in the retina (*Ttc21b* KO) and roof plate (*Gli2* KO).

Third, although we performed more detailed *in silico* analyses of selected mutants and phenotypes, we were not able to explore all mutants in detail, nor to thoroughly investigate other aspects of the data (*e.g.* the differences between wildtype strains). Even for these 22 mutants, but also looking to the future, we anticipate the community input and domain expertise will be essential to extract full value from these data, including the development of additional analytical strategies. To facilitate this, we created an interactive browser that allows exploration of mutant-specific effects on gene expression in trajectories and sub-trajectories, together with the underlying data (https://atlas.gs.washington.edu/mmca_v2/).

In 2011, the International Mouse Phenotyping Consortium (IMPC) set out to drive towards the “functionalization” of every protein-coding gene in the mouse, by generating thousands of knockout mouse lines^92^. Although over 7,000 lines have already been analysed, thousands more still await phenotyping, and even what phenotyping has been done is not necessarily comprehensive^93^. In principle, the whole embryo sc-RNA-seq phenotyping approach presented here could be extended to all Mendelian genes or even to all 20,000 mouse gene KOs, to advance our understanding of the molecular and cellular basis of human developmental disorders, to decompose pleiotropy, and to shed light on the function(s) of mammalian genes.

## Methods

### Data reporting

No statistical methods were used to predetermine sample size. Embryos used in experiments were randomised before sample preparation. Investigators were blinded to group allocation during data collection and analysis. Embryo collection and sci-RNA-seq3 analysis were performed by different researchers in different locations.

### Embryo collection

Mutants were generated through conventional gene editing tools and breeding or tetraploid aggregation and collected at the embryonic stage E13.5, calculated from the day of vaginal plug (noon = E0.5). Collection and whole embryo dissection was performed as previously described^94^. The embryos were immediately snap-frozen in liquid nitrogen and shipped to the Shendure Lab (University of Washington) in dry ice. Sets of animals with the same genotype were either all male or half male-half female. All animal procedures were in accordance with institutional, state, and government regulations.

### Nuclei isolation and fixation

Snap frozen embryos were processed as previously described^17^. Briefly, the frozen embryos were cut into small pieces with a blade and further dissected by resuspension in 1 ml ice cold cell lysis buffer (CLB, 10 mM Tris-HCl, pH 7.4, 10 mM NaCl, 3 mM MgCl2, 0.1% IGEPAL CA-630, 1% SUPERase In and 1% BSA) in a 6 cm dish. adding another 3ml CLB, the sample was strained (40 µm) into a 15 ml Falcon tube and centrifuged to a pellet (500g, 5 min). Resuspending the sample with another 1 ml CLB, the isolation of nuclei was ensured. Pelleting the isolated nuclei again (500g, 5 min) was followed by a washing step by fixation in 10 ml 4% Paraformaldehyde (PFA) for 15 minutes on ice. The fixed nuclei were pelleted (500g, 3 min) and washed twice in the nuclei suspension buffer (NSB) (500g, 5 min). The nuclei finally were resuspended in 500µl NSB and split into 2 tubes, each containing 250 µl sample. The tubes were flash frozen in liquid nitrogen and stored in a -80°C freezer, until further use for library preparation. The embryo preparation was preceded randomly for nuclei isolation in order to avoid batch effects.

### sci-RNA-seq3 library preparation and sequencing

The library preparation was performed previously described^17,95^. In short, the fixed nuclei were permeabilized, sonicated and washed. Nuclei from each mouse embryo were then distributed into several individual wells into 4 96-well plates. We split samples into four batches (∼25 samples randomly selected in each batch) for sci-RNA-seq3 processing. The ID of the reverse transcription well was linked to the respective embryo for downstream analysis. In a first step the nuclei were then mixed with oligo-dT primers and dNTP mix, denatured and placed on ice, afterwards they were proceeded for reverse transcription including a gradient incubation step. After reverse transcription, the nuclei from all wells were pooled with the nuclei dilution buffer (10 mM Tris-HCl,

pH 7.4, 10 mM NaCl, 3 mM MgCl2, 1% SUPERase In and 1% BSA), spun down and redistributed into 96-well plates containing the reaction mix for ligation. The ligation proceeded for 10 min at 25°C. Afterwards, nuclei again were pooled with nuclei suspension buffer, spun down and washed and filtered. Next, the nuclei were counted and recistributed for second strand synthesis, which was carried out at 16°C for 3h. Afterwards tagmentation mix was added to each well and tagmentation was carried out for 5 minutes at 55°C. To stop the reaction, DNA binding buffer was added and the sample was incubated for another 5 minutes. Following an elution step using AMPure XP beads and elution mix, the samples were subjected to PCR amplification to generate sequencing libraries.

Finally after PCR amplification, the resulting amplicons were pooled and purified using AMPure XP beads. The library was analysed by electrophoresis and the concentration was calculated using Qubit (Invitrogen). The library was sequenced on the NovaSeq platform (Illumina) (read 1: 34 cycles, read 2: 100 cycles, index 1: 10 cycles, index 2: 10 cycles).

### Processing of sequencing reads

Read alignment and cell-x-gene expression count matrix generation was performed based on the pipeline that we developed for sci-RNA-seq3^17^ with the following minor modifications: base calls were converted to fastq format using Illumina’s *bcl2fastq*/v2.20 and demultiplexed based on PCR i5 and i7 barcodes using maximum likelihood demultiplexing package *deML*^96^ with default settings. Downstream sequence processing and cell-x-gene expression count matrix generation were similar to sci-RNA-seq^97^ except that the RT index was combined with hairpin adaptor index, and thus the mapped reads were split into constituent cellular indices by demultiplexing reads using both the the RT index and ligation index (Levenshtein edit distance (ED) < 2, including insertions and deletions). Briefly, demultiplexed reads were filtered based on the RT index and ligation index (ED < 2, including insertions and deletions) and adaptor-clipped using *trim_galore*/v0.6.5 with default settings. Trimmed reads were mapped to the mouse reference genome (mm10), using *STAR*/v2.6.1d^98^ with default settings and gene annotations (GENCODE VM12 for mouse). Uniquely mapping reads were extracted, and duplicates were removed using the unique molecular identifier (UMI) sequence (ED < 2, including insertions and deletions), reverse transcription (RT) index, hairpin ligation adaptor index and read 2 end-coordinate (*i.e.* reads with UMI sequence less than 2 edit distance, RT index, ligation adaptor index and tagmentation site were considered duplicates). Finally, mapped reads were split into constituent cellular indices by further demultiplexing reads using the RT index and ligation hairpin (ED < 2, including insertions and deletions). To generate the cell-x-gene expression count matrix, we calculated the number of strand-specific UMIs for each cell mapping to the exonic and intronic regions of each gene with *python*/v2.7.13 *HTseq* package^99^. For multi-mapped reads, reads were assigned to the closest gene, except in cases where another intersected gene fell within 100 bp to the end of the closest gene, in which case the read was discarded. For most analyses, we included both expected-strand intronic and exonic UMIs in the cell-x-gene expression count matrix.

The single cell gene count matrix included 1,941,605 cells after cells with low quality (UMI <= 250 or detected gene <= 100) were filtered out. Each cell was assigned to its original mouse embryo on the basis of the reverse transcription barcode. We applied three strategies to detect potential doublet cells. As the first strategy, we split the dataset into subsets for each individual, and then applied the *scrublet*/v0.1 pipeline^100^ to each subset with parameters (min_count = 3, min_cells = 3, vscore_percentile = 85, n_pc = 30, expected_doublet_rate = 0.06, sim_doublet_ratio = 2, n_neighbors = 30, scaling_method = ’log’) for doublet score calculation. Cells with doublet scores over 0.2 were annotated as detected doublets (5.5% in the whole data set).

As the second strategy, we used an iterative clustering strategy based on *Seurat*/v3^101^ to detect the doublet-derived subclusters for cells. Briefly, gene count mapping to sex chromosomes was removed before clustering and dimensionality reduction, and then genes with no count were filtered out and each cell was normalized by the total UMI count per cell. The top 1,000 genes with the highest variance were selected. The data was log transformed after adding a pseudo count, and scaled to unit variance and zero mean. The dimensionality of the data was reduced by PCA (30 components) first and then with UMAP, followed by Louvain clustering performed on the 10 principal components (resolution = 1.2). For Louvain clustering, we first fitted the top 10 PCs to compute a neighbourhood graph of observations (k.param = 50) followed by clustering the cells into sub-groups using the Louvain algorithm. For UMAP visualisation, we directly fit the PCA matrix with min_distance = 0.1. For subcluster identification, we selected cells in each major cell type and applied PCA, UMAP, Louvain clustering similarly to the major cluster analysis. Subclusters with a detected doublet ratio (by *Scrublet*) over 15% were annotated as doublet-derived subclusters.

We found the above *Scrublet* and iterative clustering-based approach is limited in marking cell doublets between abundant cell clusters and rare cell clusters (*e.g.* less than 1% of the total cell population), thus, we applied a third strategy to further detect such doublet cells. Briefly, cells labeled as doublets (by *Scrublet*) or from doublet-derived subclusters were filtered out. For each cell, we only retain protein-coding genes, lincRNA genes, and pseudogenes. Genes expressed in less than 10 cells and cells expressing less than 100 genes were further filtered out. The downstream dimension reduction and clustering analysis were done with *Monocle*/v3^17^. The dimensionality of the data was reduced by PCA (50 components) first on the top 5,000 most highly variable genes and then with UMAP (max_components = 2, n_neighbors = 50, min_dist = 0.1, metric = ’cosine’). Cell clusters were identified using the Leiden algorithm implemented in *Monocle*/v3 (resolution = 1e-06). Next, we took the cell clusters identified by *Monocle*/v3 and first computed differentially expressed genes across cell clusters with the *top_markers* function of *Monocle*/v3 (reference_cells=1000). We then selected a gene set combining the top ten gene markers for each cell cluster (filtering out genes with fraction_expressing < 0.1 and then ordering by pseudo_R2). Cells from each main cell cluster were selected for dimension reduction by PCA (10 components) first on the selected gene set of top cluster-specific gene markers, and then by UMAP (max_components = 2, n_neighbors = 50, min_dist = 0.1, metric = ’cosine’), followed by clustering identification using the Leiden algorithm implemented in *Monocle*/v3 (resolution = 1e- 04). Subclusters showing low expression of target cell cluster-specific markers and enriched expression of non-target cell cluster-specific markers were annotated as doublets derived subclusters and filtered out in visualisation and downstream analysis. Finally, after removing the potential doublet cells detected by either of the above three strategies, 1,671,270 cells were retained for further analyses.

### Whole mouse embryo analysis

As described previously^17^, each cell could be assigned to the mouse embryo from which it derived on the basis of its reverse transcription barcode. After removing doublet cells and another 25 cells which were poorly assigned to any mouse embryo, 1,671,245 cells from 103 individual mouse embryos were retained (a median of 13,468 cells per embryo). UMI counts mapping to each sample were aggregated to generate a pseudobulk RNA-seq profile for each sample. Each cell’s counts were normalised by dividing its estimated size factor, and then the data were log2- transformed after adding a pseudocount followed by performing the PCA. The normalisation and dimension reduction were done in *Monocle*/v3.

We previously used sci-RNA-seq3 to generate the MOCA dataset, which profiled ∼2 million cells derived from 61 wild-type B6 mouse embryos staged between stages E9.5 and E13.5. The cleaned dataset, including 1,331,984 high quality cells, was generated by removing cells with <400 detected UMIs as well as doublets (http://atlas.gs.washington.edu/mouse-rna). UMI counts mapping to each sample were aggregated to generate a pseudobulk RNA-seq profile for each embryo. Each cell’s counts were normalised by dividing its estimated size factor, and then the data were log2-transformed after adding a pseudocount, followed by PCA. The PCA space was retained and then the embryos from the MMCA dataset were projected onto it.

### Cell clustering and annotation

After removing doublet cells, genes expressed in less than 10 cells and cells expressing less than 100 genes were further filtered out. We also filtered out low-quality cells based on the proportion of reads mapping to the mitochondrial genome (MT%) or ribosomal genome (Ribo%) (specifically, filtering cells with MT% > 10 or Ribo% > 5). We then removed cells from two embryos that were identified as outliers based on the whole-mouse embryo analysis (embryo 41 and embryo 104). This left 1,627,857 cells (median UMI count 845; median genes detected 539) from 101 individual embryos that were retained for all subsequent analyses.

To eliminate the potential heterogeneity between samples due to different mutant types and genotype backgrounds, we sought to perform the dimensionality reduction on a subset of cells from the wildtype mice (including 15 embryos with 215,575 cells, 13.2% of all cells) followed by projecting all remaining cells, derived from the various mutant embryos, onto this same embedding. These procedures were done using *Monocle*/v3. In brief, the dimensionality of the subset of data from the wildtype mice was reduced by PCA, retaining 50 components, and all remaining cells were projected onto that PCA embedding space. Next, to mitigate potential technical biases, we combined all cells from wildtype and mutant mice and applied the *align_cds* function implemented in *Monocle*/v3, with MT%, Ribo%, and log-transformed total UMI of each cell as covariates. We took the subset of cells from wildtype mice, using their “aligned” PC features to perform UMAP (max_components = 3, n_neighbors = 50, min_dist = 0.01, metric = ’cosine’) by *uwot*/v0.1.8, followed by saving the UMAP space. Cell clusters were identified using the Louvain algorithm implemented in *Monocle*/v3 on three dimensions of UMAP features, resulting in 13 isolated major trajectories (**Fig. 1e**). We then projected all of the remaining cells from mutant mouse embryos onto the previously saved UMAP space and predicted their major-trajectory labels using a *k*-nearest neighbour (*k*-NN) heuristic. Specifically, for each mutant-derived cell, we identified its 15 nearest neighbour wildtype-derived cells in UMAP space and then assigned the major trajectory with the maximum frequency within that set of 15 neighbours as the annotation of the mutant cell. We calculated the ratio of the maximum frequency to the total as the assigned score. Of note, over 99.9% of the cells from the mutant mice had an assigned score greater than 0.8. The cell-type annotation for each major trajectory was based on expression of the known marker genes (**Supplementary Table 2**).

Within each major trajectory, we repeated a similar strategy, but with slightly adjusted PCA and UMAP parameters. For the major trajectories with more than 50,000 cells, we reduced the dimensionality by PCA to 50 principal components; for the other major trajectories of more than 1,000 cells, we reduced the dimensionality by PCA to 30 principal components; for the remaining major trajectories, we reduced the dimensionality by PCA to 10 principal components. UMAP was performing with max_components = 3, n_neighbors = 15, min_dist = 0.1, metric = ’cosine’. For the mesenchymal trajectory, we observed a significant separation of cells by their cell-cycle phase in the UMAP embedding. We calculated a *g2m* index and a *s* index for individual cells by aggregating the log-transformed normalised expression for marker genes of the G2M phase and the S phase and then included them in *align_cds* function along with the other factors. Applying these procedures to all of the main trajectories, we identified 64 sub trajectories in total. Similarly, after assigning each cell from the mutant mice with a sub-trajectory label, we calculated the ratio of the maximum frequency to the total as the assigned score. Of note, over 96.7% of the cells from the mutant mice had an assigned score greater than 0.8. The cell-type annotation for each sub-trajectory was also based on the expression of known marker genes (**Supplementary Table 2**).

### Identification of inter-datasets correlated major and sub trajectories using non-negative least-squares (*NNLS*) regression

To identify correlated cell trajectories between MOCA and MMCA datasets, we first calculated an aggregate expression value for each gene in each cell trajectory by summing the log-transformed normalised UMI counts of all cells of that trajectory. For consistency during the comparison to MOCA, we manually regrouped the cells from the MMCA dataset into 10 cell trajectories, by merging the olfactory sensory neuron trajectory into the neural crest (PNS neuron) trajectory, merging the myotube trajectory, the myoblast trajectory, and the cardiomyocyte trajectory into the mesenchymal trajectory, splitting the hepatocyte trajectory into the lens epithelial trajectory and the liver hepatocyte trajectory. Next, for the two datasets, we applied non-negative least squares (*NNLS*) regression to predict gene expression in a target trajectory (*T_a_*) in dataset A based on the gene expression of all trajectories (*M_b_*) in dataset B: *T_a_ = β_0a_ + β_1a_M_b_*, based on the union of the 3,000 most highly expressed genes and 3,000 most highly specific genes in the target trajectory. We then switched the roles of datasets A and B, *i.e.* predicting the gene expression of target trajectory (*T_b_*) in dataset B from the gene expression of all trajectories (*M_a_*) in dataset A: *T_b_ = β_0b_ + β_1b_M_a_*. Finally, for each trajectory *a* in dataset A and each trajectory *b* in dataset B, we combined the two correlation coefficients: *β* = 2(*β_ab_* + 0.001)(*β_ba_* + 0.001) to obtain a statistic, where high values reflect reciprocal, specific predictivity. We repeated this analysis on sub-trajectories within each major trajectories.

### Identification of significant cell composition changes in mutant mice using beta-binomial regression

A cell number matrix of all 64 developmental sub-trajectories (*rows*) and 101 embryos (*columns*) was created and the cell number were then normalised by the size factor of each column which was estimated by *estimate_size_factors* function in *Monocle*/v3. 10 sub-trajectories with a mean of cell number across individual embryo < 10 were filtered out. The beta-binomial regression was performed using the *VGAM* package of *R*, based on the model “(trajectory specific cell number, total cell number of that embryo - trajectory specific cell number) ∼ genotype”. Of note, embryos from the four different mouse strain backgrounds were analysed independently.

### Defining and calculating lochNESS

To identify local enrichments or depletions of mutant cells, we aim to define a metric for each single cell to quantify the enrichments or depletions of mutant cells in its surrounding neighbourhood. For these analyses, we consider a mutant and a pooled wildtype combining all 4 background strains in a main trajectory as a dataset. For each dataset, we define “lochNESS” as:

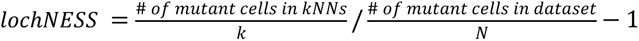

Where *N* is the total number of cells in the dataset, 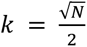 scales with N and the cells from the same embryo as the cell are excluded from the k-NNs. Note that this value is equivalent to the fold change of mutant cell percentage in the neighbourhood of a cell relative to in the whole main trajectory. For implementation, we took the aligned PCs in each sub-trajectory as calculated above and for each cell in an embryo we find the k-NNs in the remaining mutant embryo cells and wildtype cells. We plot the lochNESS in a red-white-blue scale, where white corresponds to 0 or the median lochNESS, blue corresponds to high lochNESS or enrichments, and red corresponds to low lochNESS or depletions. For reference, we simultaneously create a null distribution of lochNESS using random permutation of the mutant and wildtype cell labels, simulating datasets in which the cells are randomly mixed.

### Identifying lochNESS associated gene expression changes

To identify gene expression changes associated with mutant enriched or depleted areas, we find differentially expressed genes through fitting a regression model for each gene accounting for lochNESS. We use the *fit_models()* function implemented in *monocle/v3* with lochNESS as the *model_formula_str*. This essentially fits a generalized linear model for each gene: log(*y_i_*) = *β*_0_ + *βn* + *x_n_*., where *y_i_* is the gene expression of *gene_i_*, *β_n_* captures the effect of the lochNESS *x_n_* on expression of *gene_i_* and *β_n_* is the intercept. For each *gene_i_*, we test if *β_i_* is significantly different from zero using a Wald test and after testing all genes, we adjust the p-values using the Bejamini and Hochberg procedure to account for multiple hypotheses testing. We identify the genes that have adjusted p-value<0.05 and large positive *β_i_* values as associated with mutant enriched areas, and those with large negative *β_i_* values as associated with mutant depleted areas.

### Calculating mutant and embryo similarity scores

We can extend the lochNESS analysis, which is computed on each mutant and its corresponding wildtype mice, to compute “similarity scores” between all pairs of individual embryos from the same background strain. We consider all embryos in the same background in a main trajectory as a dataset. For each dataset, we take define a “similarity score” between *cell_n_* and *embryc_j_* as:

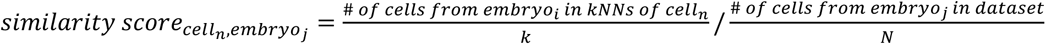

Where *N* is the total number of cells in the dataset and 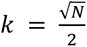. We take the mean of the similarity scores across all cells in the same embryo, resulting in an embryo similarity score matrix where entries are:

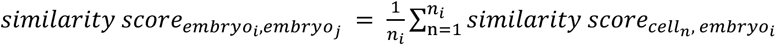

Where *n_i_* is the number of cells in *embryo_i_*. The embryo similarity score matrix can be visualised in a square heatmap where rows and/or columns are hierarchically clustered.

### Identifying and quantifying developmental delay

To identify potential mutant related developmental delay, we integrate MMCA with MOCA. We consider a mutant and its corresponding wildtype in a sub trajectory as a dataset. We take the cells from E11.5-E13.5 with similar annotations from MOCA and co-embed with the MMCA cells. We take the raw counts from both datasets, normalise, and process the data together without explicit batch correction as both datasets were generated with sci-RNA-seq3 and were similar in dataset quality. We visualise the co-embedded data in 3D UMAP space and check for developmental delay in the mutant cells (*i.e.* mutant cells embedded closer to early MOCA cells compared to wildtype cells). To quantify the amount of developmental delay, we find k-NNs in MOCA for each cell in MMCA and calculate 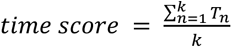, where *T_n_* is the developmental time of MOCA *cell_n_* in the k-NNs of the MMCA cell. Afterwards, we test if the average time scores of mutant cells are significantly different from that of wildtype cells using a student’s t-test.

### RNAscope *in situ* Hybridization

For RNAscope, embryos were collected at stage E13.5 and fixed for 4 hours in 4% PFA/PBS at room temperature. The embryos were washed twice in PBS before incubation in a sucrose series (5%, 10% and finally 15% sucrose (Roth) /PBS) each for an hour or until the embryos sank to the bottom of the tube. Finally, the embryos were incubated in 15% sucrose/PBS and O.C.T. (Sakura) in a 1:1 solution before embedding the embryos in O.C.T in a chilled ethanol bath and put into - 80°C for sectioning. The embryos were cut into 5 μm thick sections on slides for RNAscope.

Simultaneous RNA *in situ* hybridization was performed using the RNAscope® technology (Advanced Cell Diagnostics [ACD]) and the following probes specific for Mm-Kcnj2 (Cat. No. 476261, ACD) and Mm-Sox9 (Cat. No. 401051-C2, ACD) on five μm sections of the mouse embryos. RNAscope probes were purchased by ACD and designed as described by Wang *et al.*^102^. The RNAscope® assay was run on a HybEZ™II Hybridization System (Cat. No. 321720, ACD) using the RNAscope® Multiplex Fluorescent Reagent Kit v2 (Cat. No. 323100, ACD) and the manufacturer’s protocol for fixed-frozen tissue samples with target retrieval on a hotplate for 5 minutes. Fluorescent labelling of the RNAscope® probes was achieved by using OPAL 520 and OPAL 570 dyes (Cat. No. FP1487001KT + Cat. No. FP1488001KT, Akoya Biosciences, Marlborough, MA, USA) and stained sections were scanned at 25x magnification using a LSM 980 with Airyscan 2 (Carl Zeiss AG, Oberkochen, DE).

### Image analysis

For quantitative analysis of the RNAscope images, representative fields of view for each stained section were analysed using the image processing software Fiji^103^. Each organ of interest mRNA signal was counted in a defined area (1×1 mm^2^) with an n=6 per condition. Statistics were calculated using student t-Test and evaluated (- p > 0,05 = non-significant, p < 0,05 - ≥ 0,01 = *, p < 0,01 - ≥ 0,001= ** - p < 0,001= ***).

### Clustering and annotation limb mesenchyme trajectory

*Seurat/v4.0.6* was used for the analysis. Wildtype cells in the limb mesenchyme trajectory from all wild-type mice (n = 15 mice, n = 25,211 cells) were used to first annotate the cells. The raw counts were log-normalised after which PCA was performed with default parameters on top 2000 highly variable genes selected using the “vst” method. Nearest neighbours were computed on the PCA space, with default parameters, except that all the principal components computed earlier were used. Clustering was performed using the Louvain community detection algorithm with a resolution of 0.1, resulting in three clusters. Positive marker genes for these clusters were identified using the Wilcoxon Rank Sum test, where only the genes expressed in at least 20% of the cells in either cell groups were considered. The clusters were annotated based on biologically relevant markers (**Supplementary Fig. 12b**). The newly assigned cell annotations for the Limb mesenchyme trajectory cells in the wildtype dataset were transferred to the corresponding cells in the S*ox9* regulatory INV mutant using the *FindTransferAnchors* and *TransferData* functions using default parameters, except that all the computed principal components were used. 92.3% of the transferred annotations had a score (prediction.score.max) greater than or equal to 0.8.

### Density visualization and RNA velocity analysis

Using *Seurat/v4.0.6*, the raw counts were log-normalised, and PCA was performed with default parameters on top highly variable genes 2000 genes, selected using the “*vst*” method. Dimensionality reduction was performed using PCA using default parameters, after which the UMAP embedding was carried out on all computed PC components. Density plots were created using the *stat_2d_density_filled* function in *ggplot2/v3.3.5*. For RNA velocity analysis using *scVelo/v0.2.4*, the total, spliced, and unspliced count matrices, along with the UMAP embeddings were exported as an h5ad file using *anndata/v0.7.5.2* for R. The count matrices were filtered and normalised using *scv.pp.filter_and_normalize*, with *min_shared_counts*=20 and *n_top_genes*=2000. Means and variances between 30 nearest neighbours were calculated in the PCA space (*n_pcs*=50, to be consistent with default value in *Seurat*). The velocities were calculated using default parameters and projected onto the UMAP embedding exported from *Seurat*.

### Single sample Gene Set Enrichment Analysis

Single-sample Gene Set Enrichment Analysis (ssGSEA) was applied to sc-RNA-seq data using the *escape* R-package^84^. The *msigdbr* and *getGeneSets* functions were used to fetch and filter the entire Hallmark (H, 50 sets) or the Signature Cell Type (C8, 700 sets) *Mus musculus* gene sets from the MSigDB^104,105^. *enrichIt* with default parameters, except for using 10000 groups and variable number of cores, was performed on the seurat-object containing data corresponding to the undifferentiated mesenchyme cells from the *Sox9* regulatory INV mutant, after converting the feature names to gene symbols as necessitated by the *escape* package. The obtained enrichment scores for each gene set were compared between the two branches (**Fig. 4f**) using the two sample Wilcoxon test (*wilcox_test*) with default parameters and adjusted for multiple comparisons using Bonferroni correction.

## Data availability

The data generated in this study can be downloaded in raw and processed forms from the NCBI Gene Expression Omnibus under accession number GSE199308. Other intermediate data files, code and an interactive app to explore our dataset will be made freely available via https://atlas.gs.washington.edu/mmca_v2/.

## Code availability

All code will be made freely available through a public GitHub repository.

## Acknowledgements

We thank Stefan Mundlos and Cesar Prada for helpful discussions around data processing and analysis and results interpretation, as well as all members of the Cao, Shendure, Spielmann labs for continuous support and helpful input. N.Haag and I.K. thank Matthias Ebbinghaus for help with breeding of Scn11a GOF mice. We thank Vanessa Suckow for genotyping the Ror2 KI and Cdkl5 -/Y mice. We thank Scott Houghtaling and Tzu-Hua Ho for breeding and embryo harvest of *Ttc21b, Carm1* and *Gli2* mice. X.H. thanks Gwen the Cat for support and cheer ups during meetings. J.S. and work in the Shendure Lab was supported by the Paul G. Allen Frontiers Foundation (Allen Discovery Center grant to J.S. and C.T.), the National Institutes of Health (grant UM1HG011531 to J.S.), Alex’s Lemonade Stand’s Crazy 8 Initiative (to J.S.) and the Bonita and David Brewer Fellowship (C.Q.). Work at the E.O. Lawrence Berkeley National Laboratory was supported by U.S. National Institutes of Health (NIH) grants to L.A.P. and A.V. (UM1HG009421 and R01HG003988) and performed under U.S. Department of Energy Contract DE-AC02- 05CH11231, University of California. J.S. is an Investigator of the Howard Hughes Medical Institute. M.S. is a DZHK principal investigator and is supported by grants from the Deutsche Forschungsgemeinschaft (DFG) (SP1532/3-1, SP1532/4-1, and SP1532/5-1) and the Deutsches Zentrum für Luft- und Raumfahrt (DLR 01GM1925). D.R.B was supported by R01HD36404 from NICHD. J.C. is supported by the National Institutes of Health (grant DP2 HG012522-01 and RM1HG011014) and the Rockefeller University.

## Author contributions

J.C., M.S. and J.S. conceptualized, supervised and funded the project. D.R.B., W.C., A.D., D.E.D., N.Haag, D.I., I.K., F.H., V.M.K., U.K., L.A.P., S.R., A.R., M.R., A.V. L.W. and Y.Z. provided mouse embryos. J.C. and J.H. extracted and fixed the nuclei from embryos and performed the sci-RNA-seq experiment. S.U., R.B., R.H., N.Hans. and J.H. performed RNAscope experiment and image analysis. X.H., C.Q., J.H., V.S. and S.B. performed all computational analyses. C.M. created the interactive webpage with guidance from X.H. and J.S. L.S., S.S. and C.T. provided assistance with data analysis and results interpretation. X.H., C.Q., J.H. and V.S. wrote the first draft of the manuscript, which was finalized together with J.C., M.S. and J.S. and input from all authors.

## Competing interests

J.S. is a SAB member, consultant and/or co-founder of Cajal Neuroscience, Guardant Health, Maze Therapeutics, Camp4 Therapeutics, Phase Genomics, Adaptive Biotechnologies and Scale Biosciences.

---

**Supplementary Figure 1.**
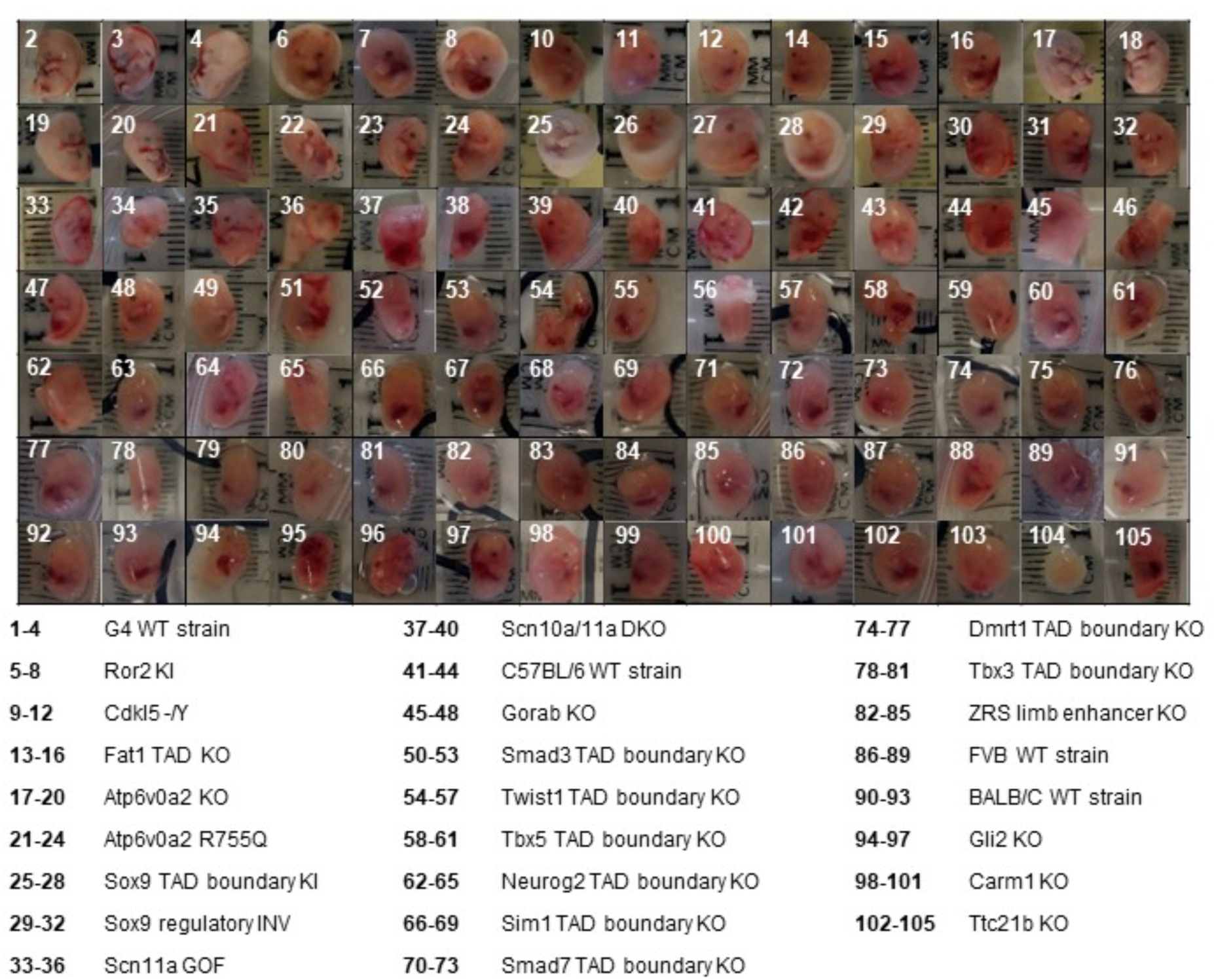
Images of mouse embryos. 104 embryos (26 genotypes x 4 replicates) were staged at E13.5 and sent by five groups to a single site. #49 was accidentally skipped in our numbering systems. Embryo #70 was lost in transport. Pictures of embryos #1, #5, #9, #13 and #91 were not taken, but the embryos were included in the sci-RNA-seq3 experiment. As discussed in the text, embryos #41 and #104 were labelled as outliers based on computational analyses and their data discarded, while data from the remaining 101 embryos were retained and analysed further. Of note, in addition to the computational analyses suggesting that embryo #104 was an outlier, it was also relatively small in size upon visualisation.

**Supplementary Figure 2.**
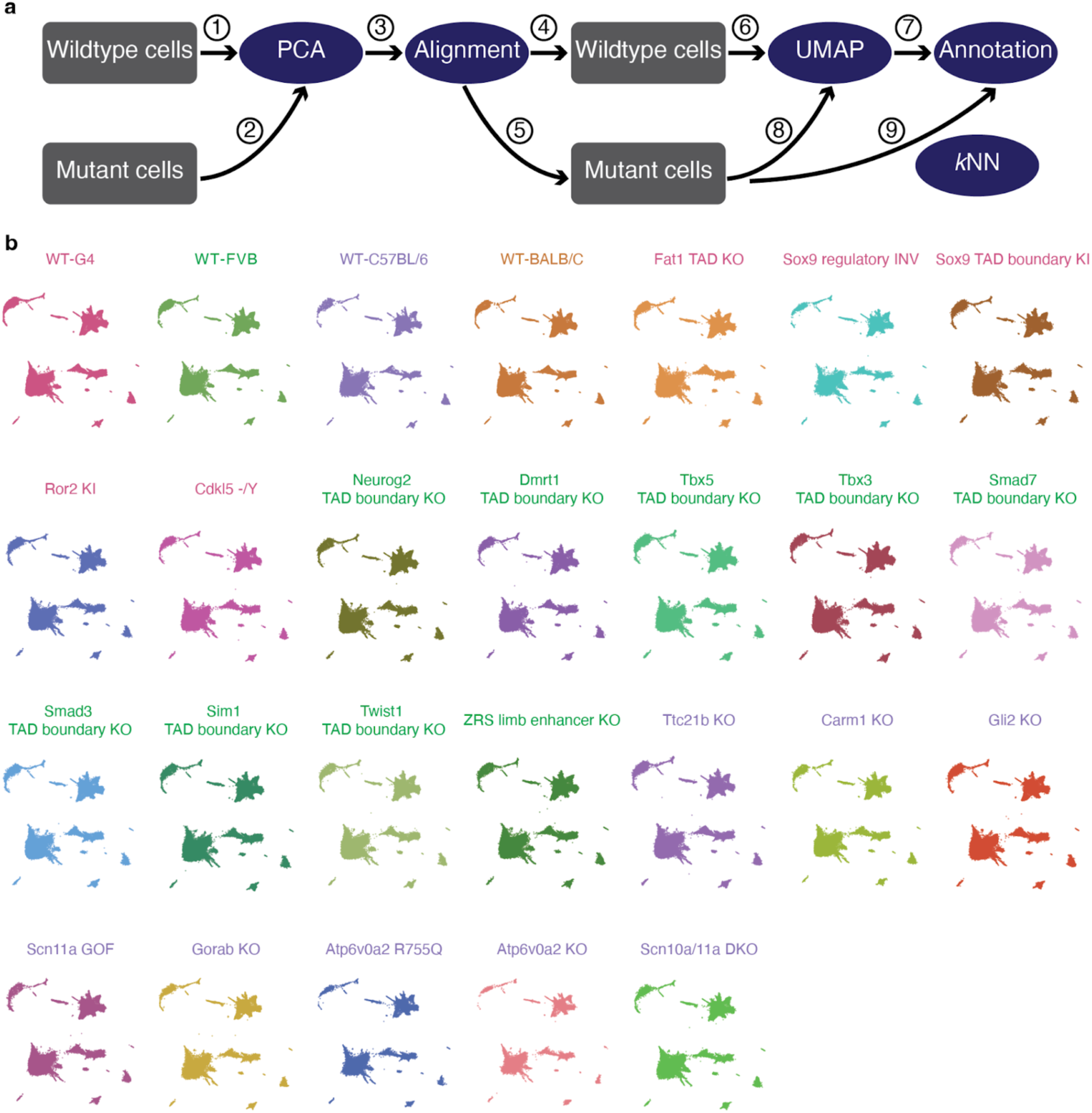
Integrating cells derived from embryos of multiple genetic backgrounds to a single, wildtype-based “reference embedding”. **a,** Schematic of approach. We first applied principal components-based dimensionality reduction to cells from wildtype genotypes only (①). We then projected cells from the mutant embryos to this PCA embedding (②). Next, to mitigate potential biases from technical factors, we applied the *align_cds* function in *Monocle*/v3, with the MT%, Ribo%, and log-transformed total UMIs of each cell as covariates (③). We then split wildtype and mutant cells again (④ & ⑤), and applied the UMAP algorithm to wildtype cells only using their “aligned” PC features (⑥), followed by Louvain clustering and manual annotation of individual clusters based on marker gene expression to identify major trajectories, and then iterative clustering and annotation to identity and annotate sub-trajectories (⑦). Finally, cells from mutant embryos were projected to this wildtype-based UMAP embedding, again using their aligned PC features (⑧). Major trajectory labels were assigned to mutant cells via a *k*-nearest neighbour (*k*-NN) heuristic, and these last steps were repeated to further assign sub-trajectory labels to mutant cells (⑨). **b,** 3D UMAP visualisations of cells from each wildtype or mutant background within the shared “reference embedding” resulting from the aforedescribed procedures.

**Supplementary Figure 3.**
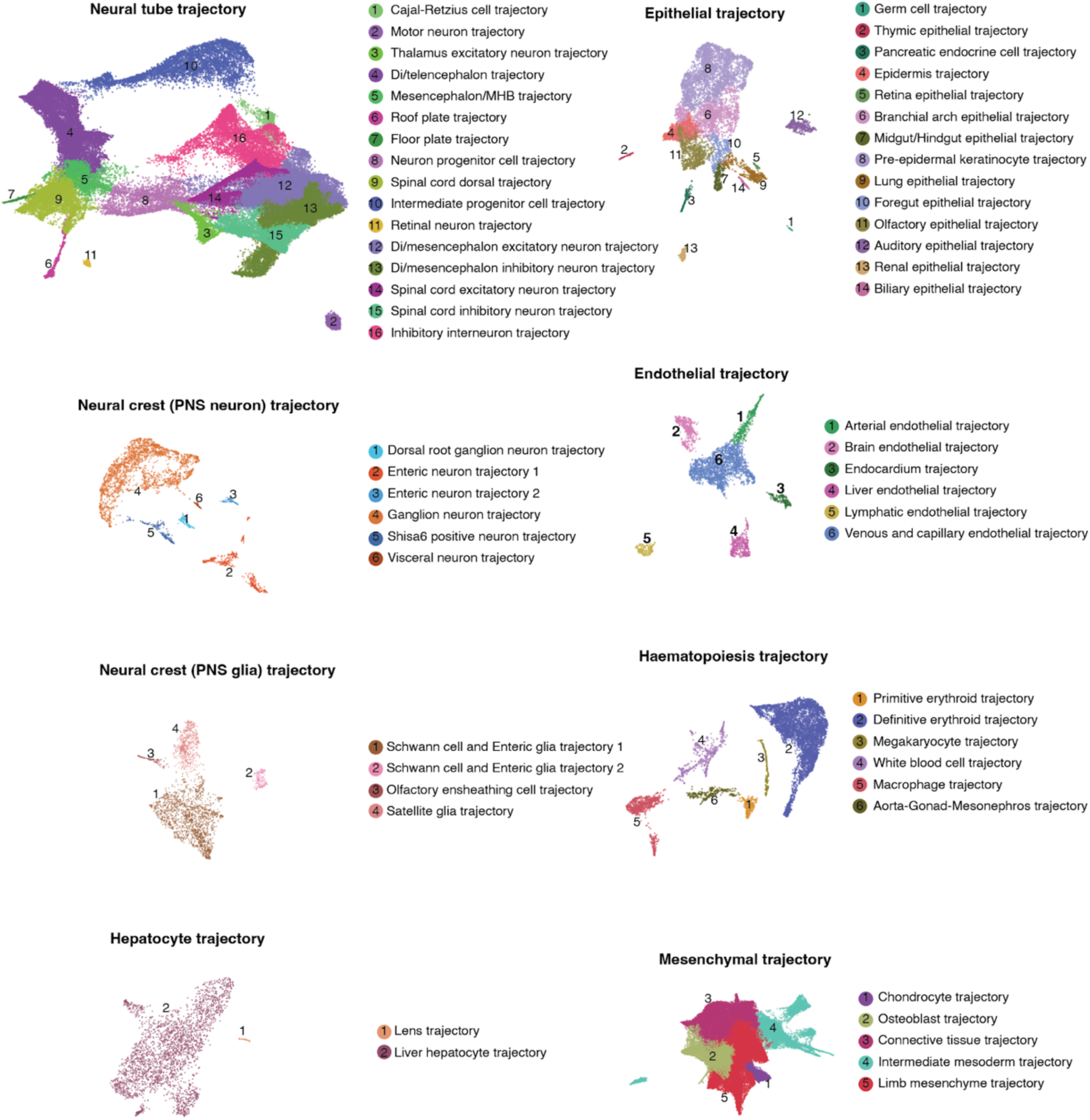
Annotation of sub-trajectories in data from wildtype E13.5 embryos. From 215,517 single cell profiles of wildtype E13.5 embryos of four strains in MMCA, we annotated 13 major trajectories. For 8 of these 13 major trajectories, iterative analysis identified the additional sub-trajectories shown here as 3D UMAP visualisations. Cells are colored by sub-trajectory annotations. PNS: peripheral nervous system. MHB: midbrain-hindbrain boundary. Di: Diencephalon.

**Supplementary Figure 4.**
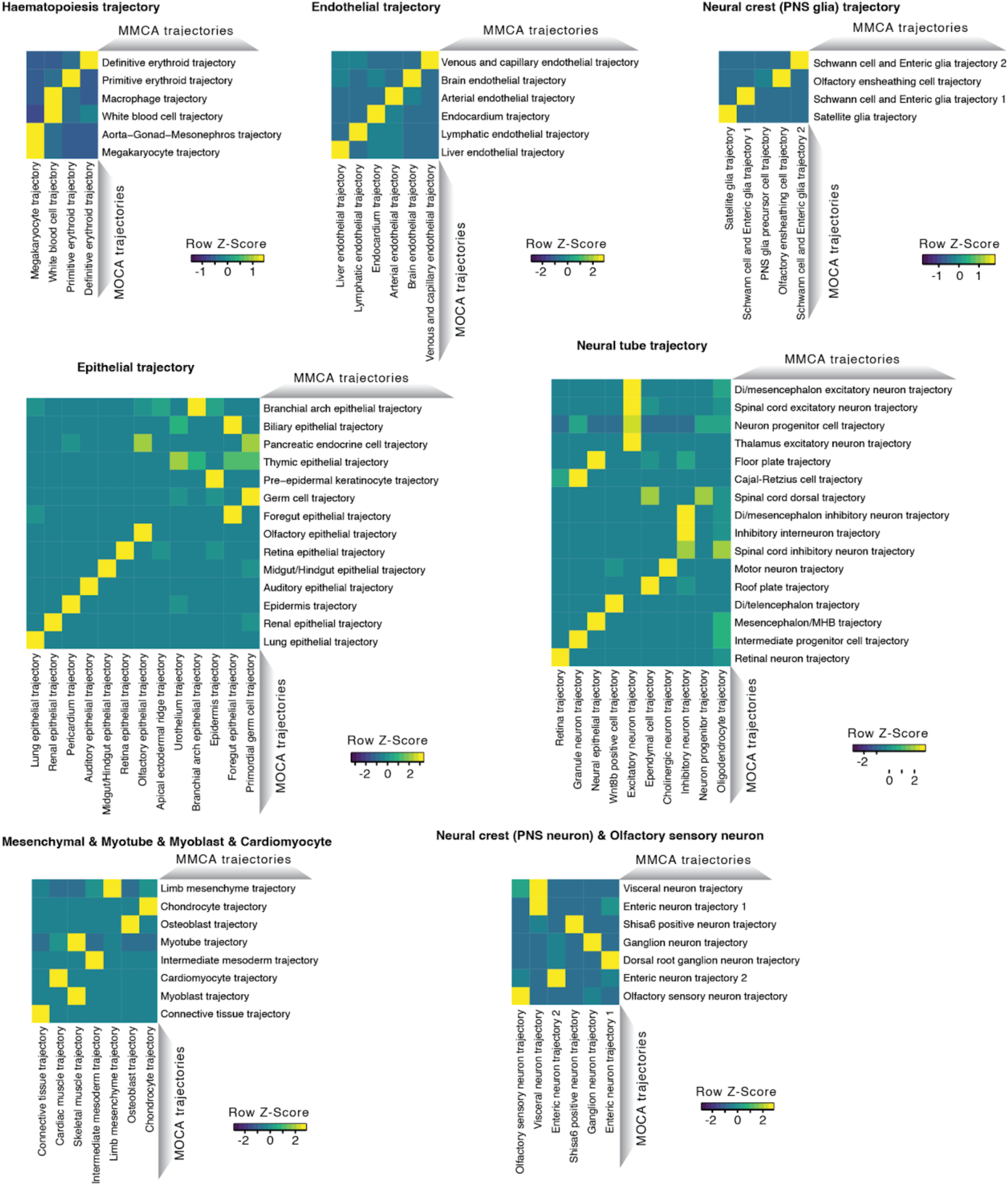
Correlated developmental sub-trajectories between MOCA (E9.5 - E13.5) and MMCA (E13.5 only) based on non-negative least-squares (*NNLS*) regression. Similar to **Fig. 1f**, shown here are heat maps of the combined *β* values (row-scaled) between developmental sub-trajectories from MMCA (rows) and developmental sub-trajectories from the MOCA (columns), within each major trajectory. PNS: peripheral nervous system. MHB: midbrain-hindbrain boundary. Di: Diencephalon.

**Supplementary Figure 5.**
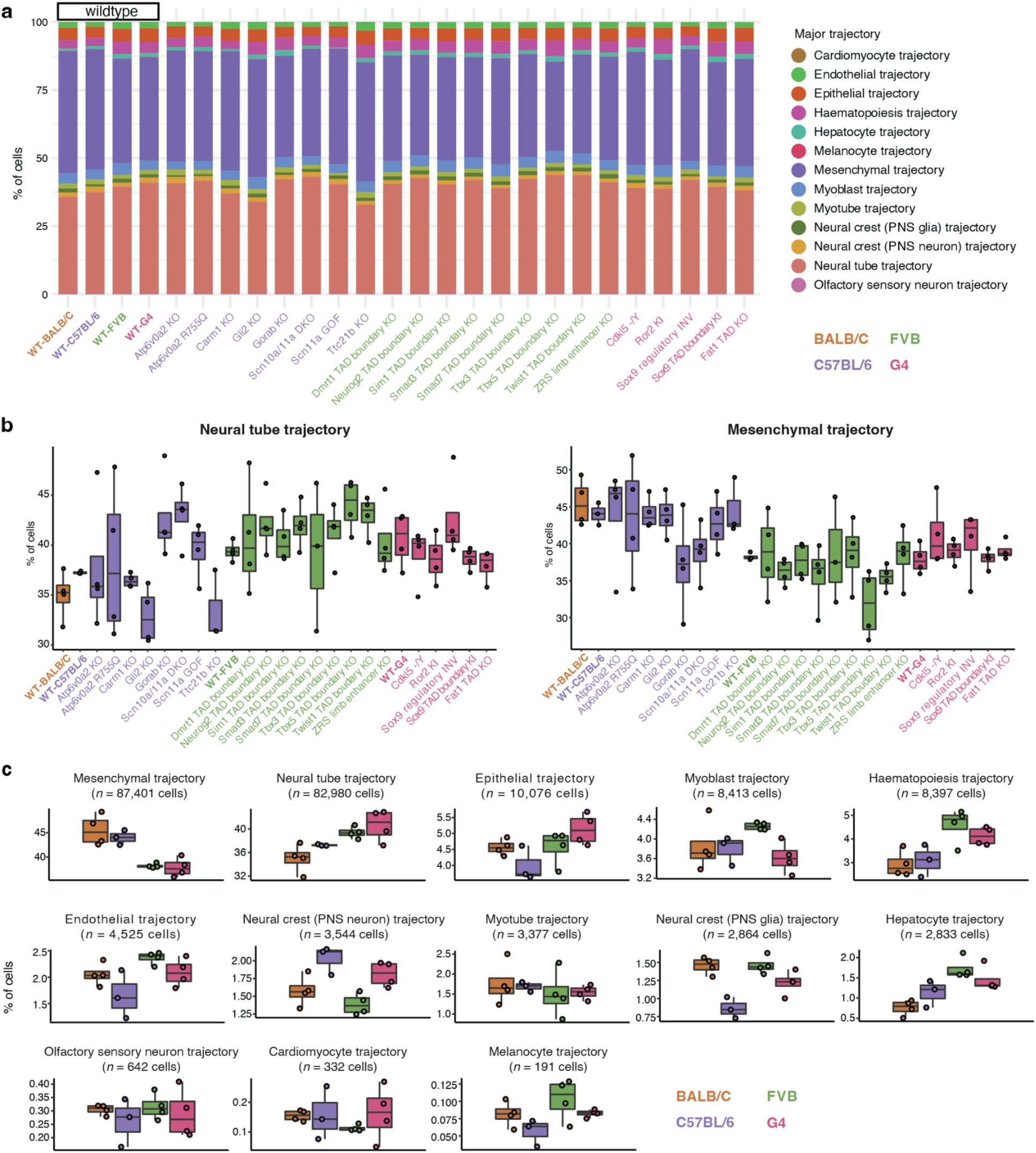
Cell composition for individual wildtype and mutant embryos across developmental trajectories. **a,** Cell composition across 13 major trajectories of embryos from different wildtype or mutant strains. Cells from all replicates for each strain were pooled for this visualisation. **b,** Boxplots of cell proportions falling into neural tube (left) or mesenchymal (right) trajectories for different wildtype or mutant strains. Each point corresponds to an individual embryo. **c,** Boxplots of cell proportions falling into each of the 13 major trajectories for the four wildtype strains. Each point corresponds to an individual embryo. The total number of cells from each major trajectory profiled from wildtype embryos is also listed. In the boxplots (panels b & c), the centre lines show the medians; the box limits indicate the 25th and 75th percentiles; the replicates are represented by the dots. PNS: peripheral nervous system.

**Supplementary Figure 6.**
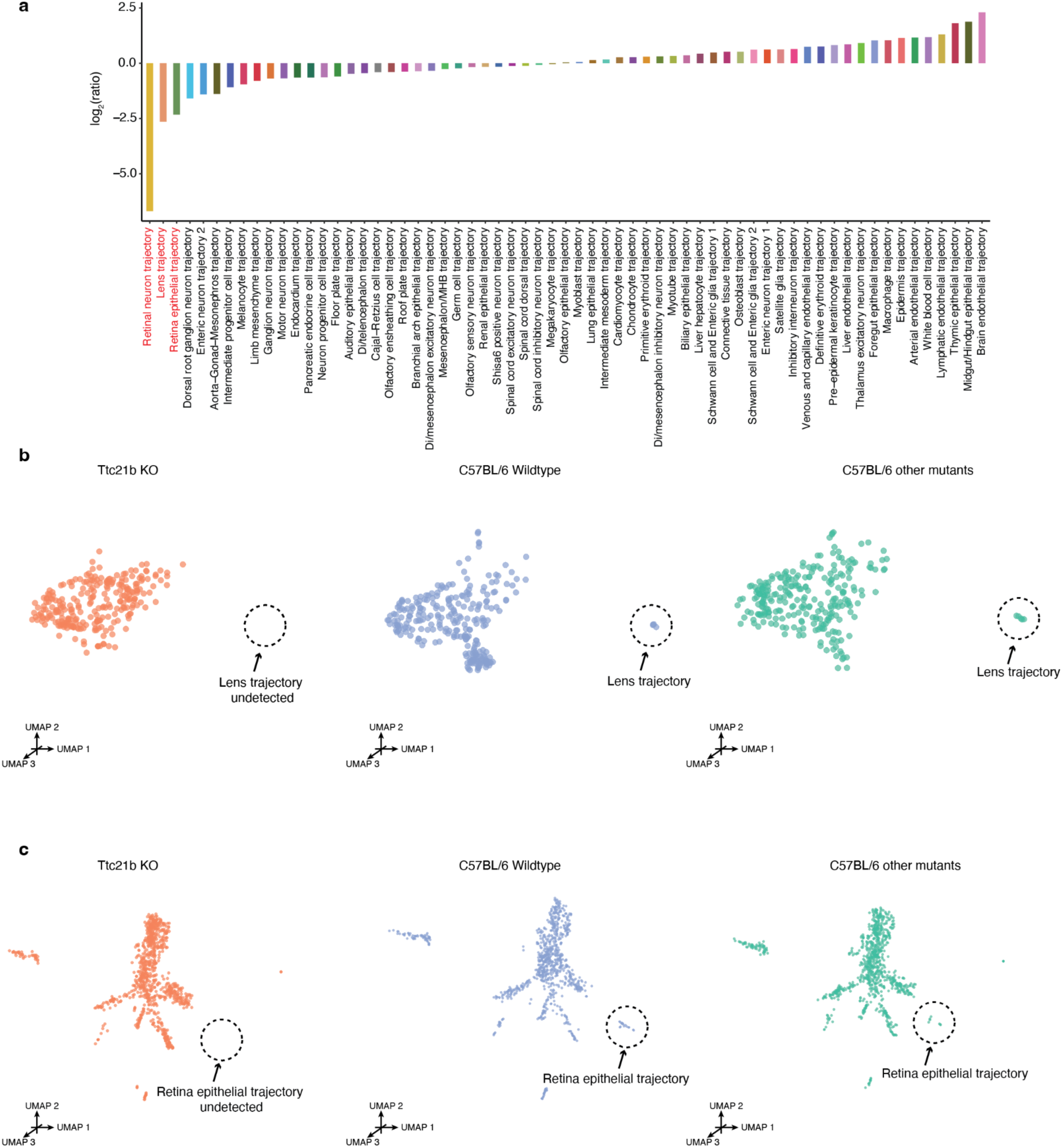
Multiple retinal trajectories are diminished in *Ttc21b* KO mice. **a,** The log2 transformed ratio of the cell proportions of each sub-trajectory, comparing *Ttc21b* KO and C57BL/6 wildtype embryos, are shown. Although reductions in the retina epithelial and lens trajectories were excluded from the regression analysis due to their low numbers, they were, together with the retinal neuron trajectory, the most extreme in magnitude. **b,** 3D UMAP visualisation of the hepatocyte major trajectory, highlighting cells from either the *Ttc21b* KO (left), C57BL/6 wildtype (middle), or other mutants on the C57BL/6 background (right). The three plots were randomly downsampled to the same number of cells (*n* = 264 cells) **c,** 3D UMAP visualisation of the epithelial major trajectory, highlighting cells from either the *Ttc21b* KO (left), C57BL/6 wildtype (middle), or other mutants on the C57BL/6 background (right). The three plots were randomly downsampled to the same number of cells (*n* = 937 cells).

**Supplementary Figure 7.**
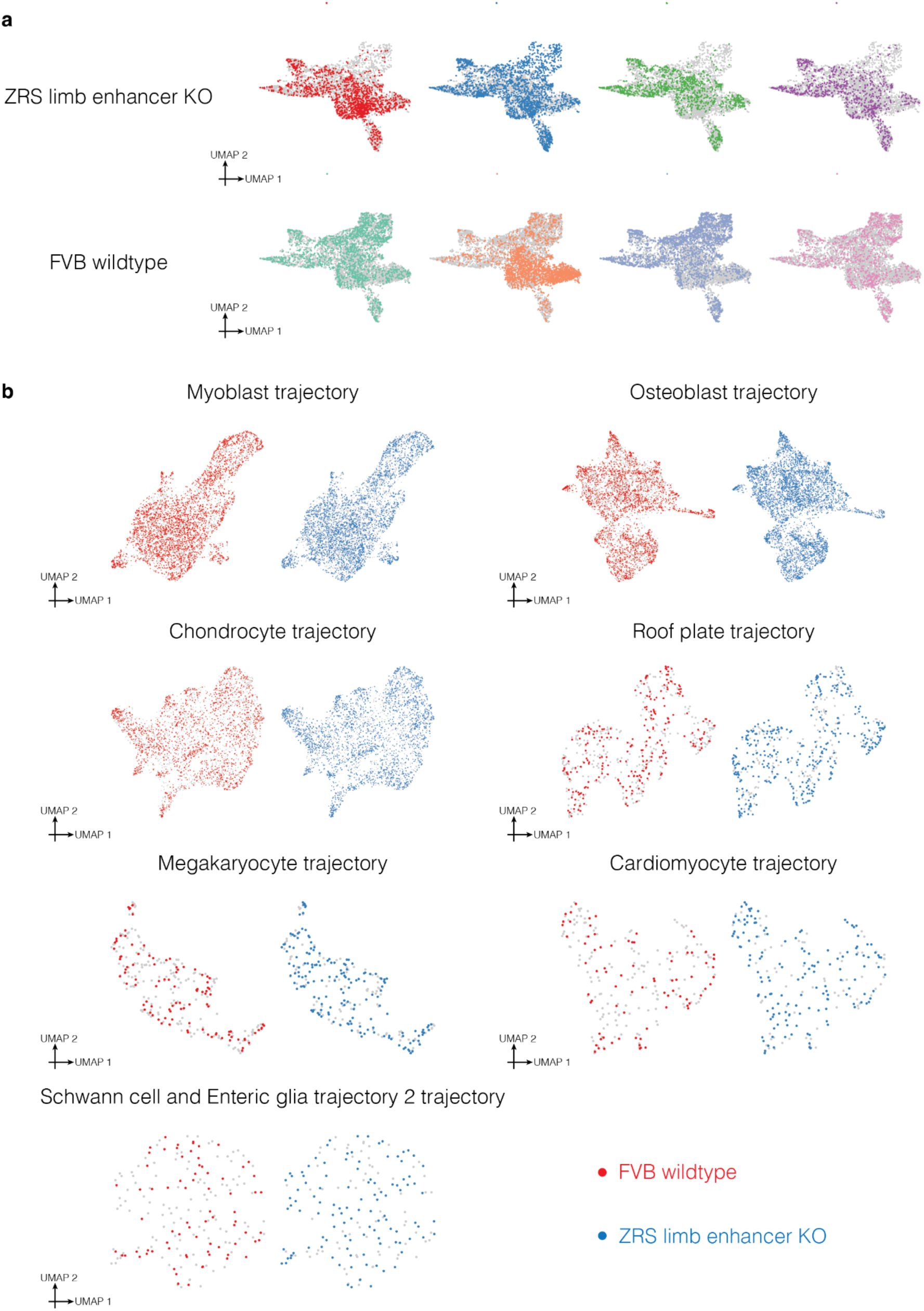
Co-embedding cells from nominally altered trajectories from ZRS limb enhancer KO and FVB wildtype. **a,** UMAP visualisation of co-embedded cells of limb mesenchyme trajectory from the ZRS limb enhancer KO and FVB wildtype. The same UMAP is shown eight times, highlighting cells from either ZRS limb enhancer KO (top row) or FVB wildtype (bottom row), and breaking out the four individual replicates for each strain. **b,** UMAP visualisation of co-embedded cells of various sub-trajectories from the ZRS limb enhancer KO and FVB wildtype. The same UMAP is shown twice for each, highlighting cells from either FVB wildtype (left) or ZRS limb enhancer KO (right). These are the seven sub-trajectories in which, in addition to limb mesenchyme, we detected nominally significant differences in cell type proportions for the ZRS limb enhancer KO.

**Supplementary Figure 8.**
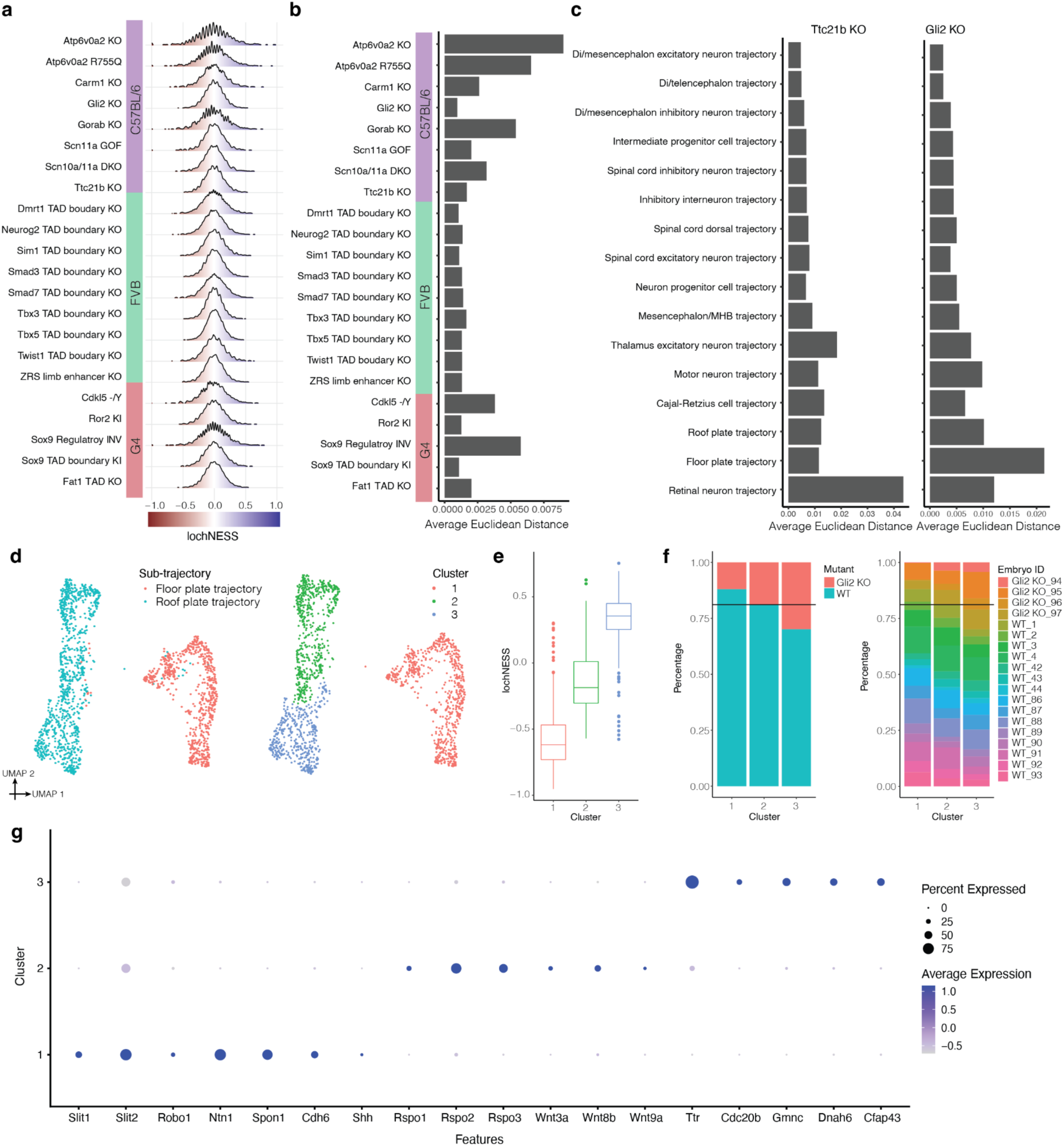
Quantitative analysis of lochNESS distributions and analysis of *Gli2* KO in the roof plate and floor plate trajectories. **a,** Distribution of lochNESS in all cells of each mutant under random permutation of mutant labels. **b,** Barplot showing the average euclidean distance between lochNESS vs. lochNESS under permutation across all cells within a mutant. **c,** Barplots showing the average euclidean distance between lochNESS and lochNESS under permutation, across all cells in neural tube sub-trajectories of the *Ttc21b* KO and *Gli2* KO mutants. **d,** UMAP visualisation of co-embedded cells of the floor plate and roof plate sub-trajectories from the *Gli2* KO mutant and pooled wildtype, colored by sub-trajectory (left) or cluster number (right). **e,** Boxplot showing the lochNESS distribution in each cluster shown on the right of panel **d**. **f,** Barplots showing the cell composition of each cluster shown on the right of panel **d**, split by mutant vs. wildtype (left) or individual embryo (right), with a reference line at the overall wildtype cell proportion. **g,** Dotplot summarising the expression of and percent of cells expressing selected marker genes in each cluster shown on the right of panel **d**.

**Supplementary Figure 9.**
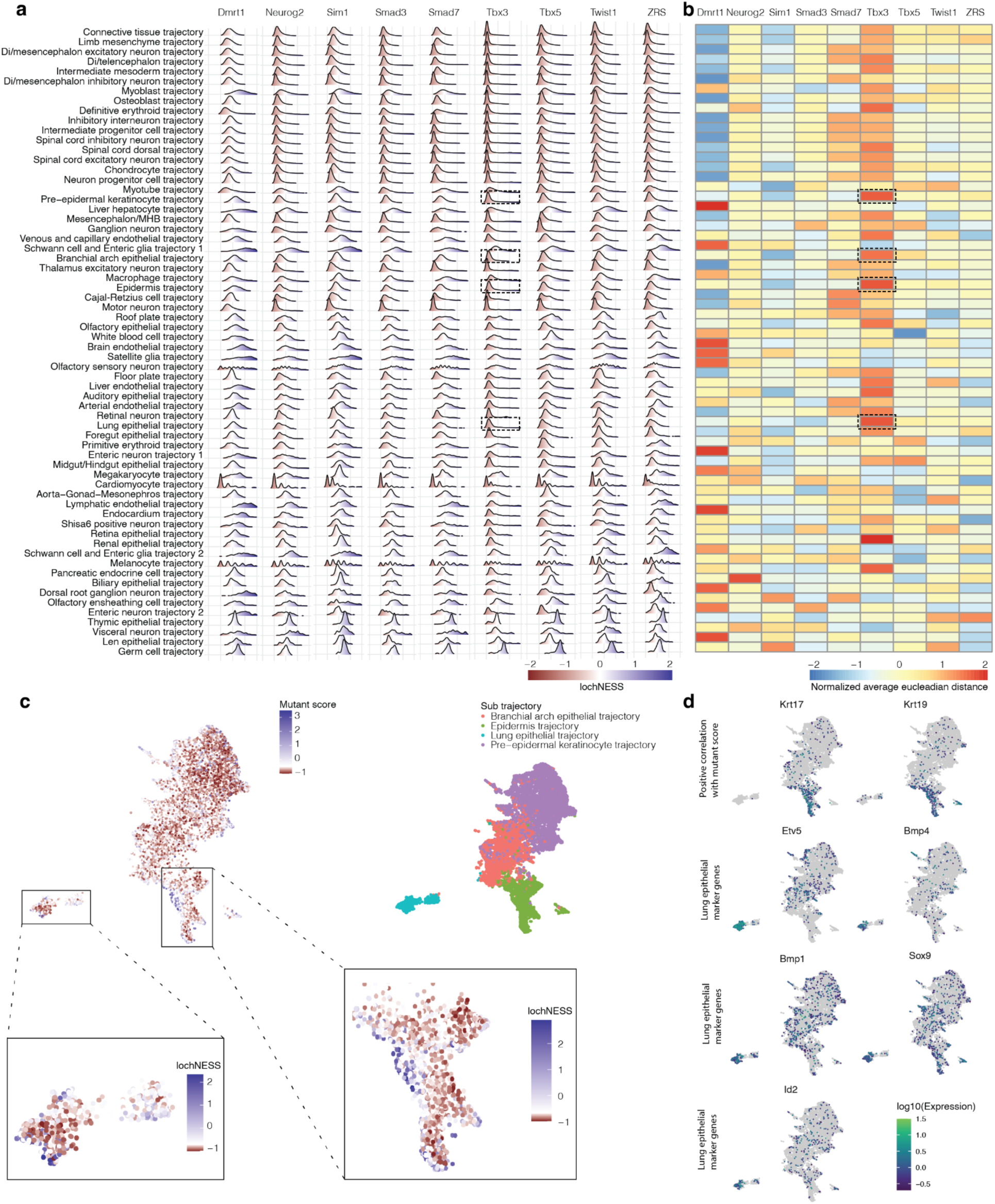
Systematic screening of lochNESS distributions identifies altered epithelial sub-trajectories in the *Tbx3* TAD Boundary KO mutant. **a,** Distribution of lochNESS in each sub-trajectory of the mutants in the FVB background strain, all of which are TAD boundary KOs. Dashed boxes in the sixth column highlight the most deviated epithelial sub-trajectories in the *Tbx3* TAD Boundary KO mutant. **b,** Row-normalised heatmap showing the average euclidean distance between lochNESS and lochNESS under permutation in each sub-trajectory for the same mutants shown in panel **a**, centred and scaled by row. Dashed boxes in the sixth column again highlight the most deviated epithelial sub-trajectories in the *Tbx3* TAD Boundary KO mutant. **c,** UMAP showing co-embedding of *Tbx3* TAD Boundary KO and pooled wildtype cells in the pre−epidermal keratinocyte, epidermis, branchial arch, and lung epithelial sub trajectories, colored by lochNESS (top left) [with blown up insets showing lochNESS in lung epithelial (bottom left) and epidermis (bottom right) sub-trajectories], or by sub-trajectory identity (right). LochNESS colour scale is centred at the median of lochNESS. **d,** same as in panel **c**, but colored by expression of selected mutant related genes and marker genes.

**Supplementary Figure 10.**
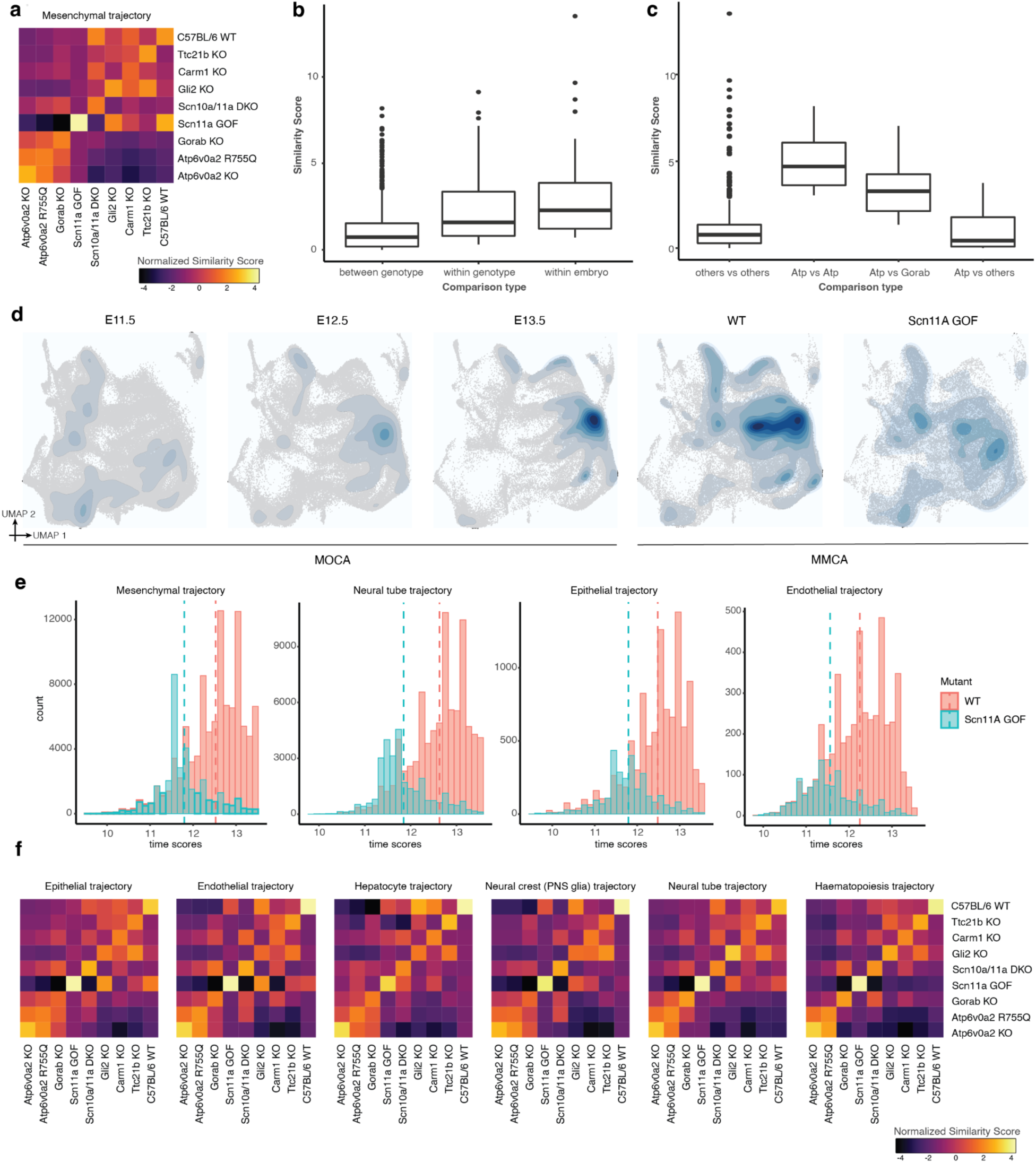
Similarity scores reveal mutant-shared and mutant-specific effects. **a,** Heatmap showing similarity scores between C57BL/6 genotypes in the mesenchymal trajectory. **b,** Boxplot showing the similarity scores of comparisons between embryos of different genotypes (left), between embryos of the same genotype (middle), and within the same embryos (right) for C57BL/6 genotypes in the mesenchymal trajectory. **c,** Boxplot showing the similarity scores of comparisons between *Atp6v0a2* KO vs. *Atp6v0a2* R755Q (left), *Atp6v0a2* KO or *Atp6v0a2* R755Q vs. *Gorab* KO (middle), *Atp6v0a2* KO or *Atp6v0a2* R755Q vs. other C57BL/6 genotypes, in the mesenchymal trajectory. Genotype names are simplified in the x-axis legend (“Atp” = *Atp6v0a2* KO or *Atp6v0a2*, “Gorab” = *Gorab* KO, “others” = *Carm1* KO, *Gli2* KO, *Scn10a/11a* DKO, *Scn11a* GOF, *Ttc21b* KO or C57BL/6 wildtype). **d,** UMAPs showing co-embedding of *Scn11a* GOF cells with pooled wildtype cells and E11.5-E13.5 MOCA cells, in the neural tube trajectory, split by mutant (MMCA) and time point (MOCA), with cell density and distributions overlaid. **e,** Barplots showing the distribution of “time scores” for *Scn11a* GOF cells and pooled wildtype cells in the mesenchyme, neural tube, endothelial and epithelial main trajectories, with reference lines at the mean value of time scores. **f,** Heatmaps showing similarity scores between C57BL/6 genotypes in selected main trajectories. *Gorab* KO exhibits high similarity to the two *Atp6v0a2* genotypes in the epithelial, endothelial, hepatocyte and neural crest (PNS glia) trajectories, but not the neural tube and hematopoiesis trajectories.

**Supplementary Figure 11.**
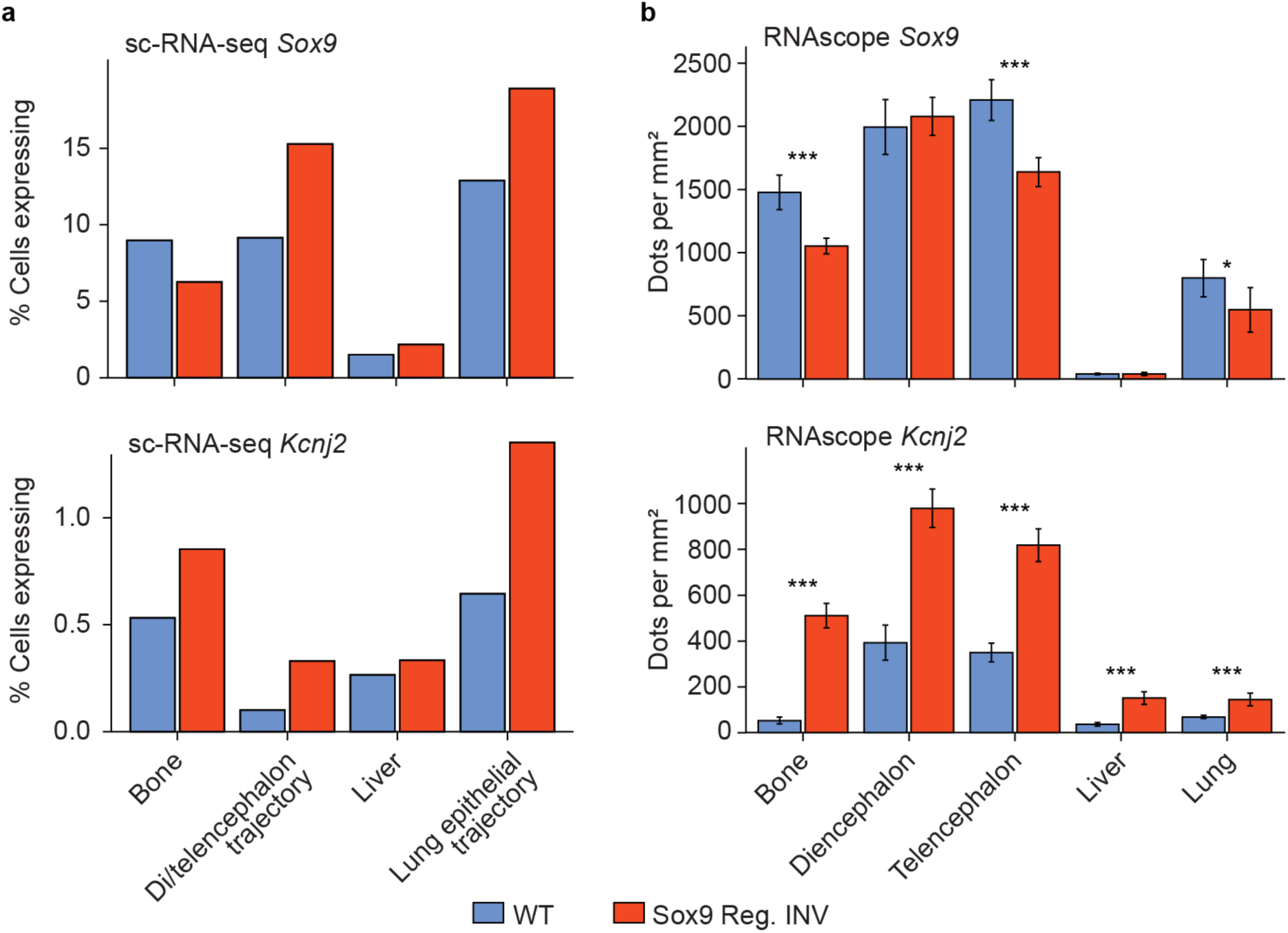
Misregulation of *Sox9* and *Kcnj2* in the *Sox9* regulatory INV mutant. **a,** Quantification of *Sox9* (top row) and *Kcnj2* (bottom row) expression in sc-RNA-seq data in the wildtype (blue) and *Sox9* regulatory INV (red) genotypes in selected trajectories. For “bone” and “liver”, multiple sub-trajectories were pooled to match the tissue labels in the RNAscope data in panel **b**. Specifically, “bone” refers to cells from chondrocyte, osteoblast, and limb mesenchyme trajectories, whereas “liver” refers to cells from the liver endothelial and liver hepatocyte trajectories. **b,** Quantification of *Sox9* and *Kcnj2* expression based on RNAscope images of heterozygous E13.5 wildtype and Sox9 regulatory INV mutant embryos (images not shown; available upon request). The mRNA signal was counted in a defined area (1×1 mm^2^), n=6 each condition. Statistics were calculated using student t-test and evaluated the following: p > 0.05 = non-significant; p < 0,05 - ≥ 0.01 = *; p < 0,01 - ≥ 0.001= **; p < 0.001= ***.

**Supplementary Figure 12.**
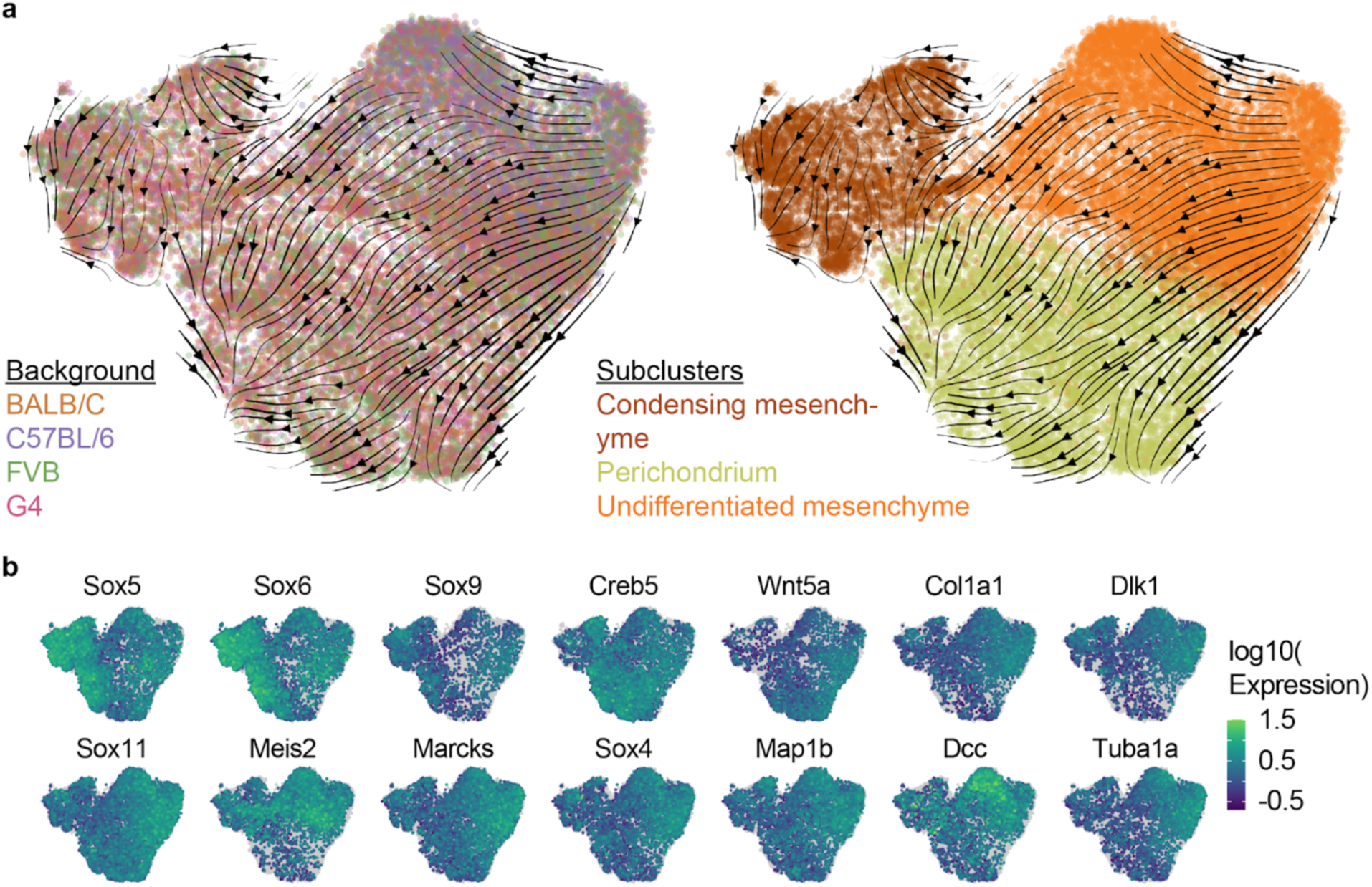
Sub-clustering and annotation of the wildtype limb mesenchyme. **a,** Sub- clustering of the limb mesenchyme trajectory based on cells from pooled wildtype. RNA velocity arrows generated using scVelo (Methods) indicate the transition of undifferentiated mesenchyme (marked by *Meis2, Marcks, Map1b*) into perichondrium (*Wnt5a,Creb5*) and condensing mesenchyme (*Sox5, Sox6, Sox9*) in all wildtype samples^86–91^. **b,** Marker gene expression used to annotate limb mesenchyme sub- clusters. All except *Dcc* and *Tuba1a* are literature-based markers of the three cell types.

**Supplementary Figure 13.**
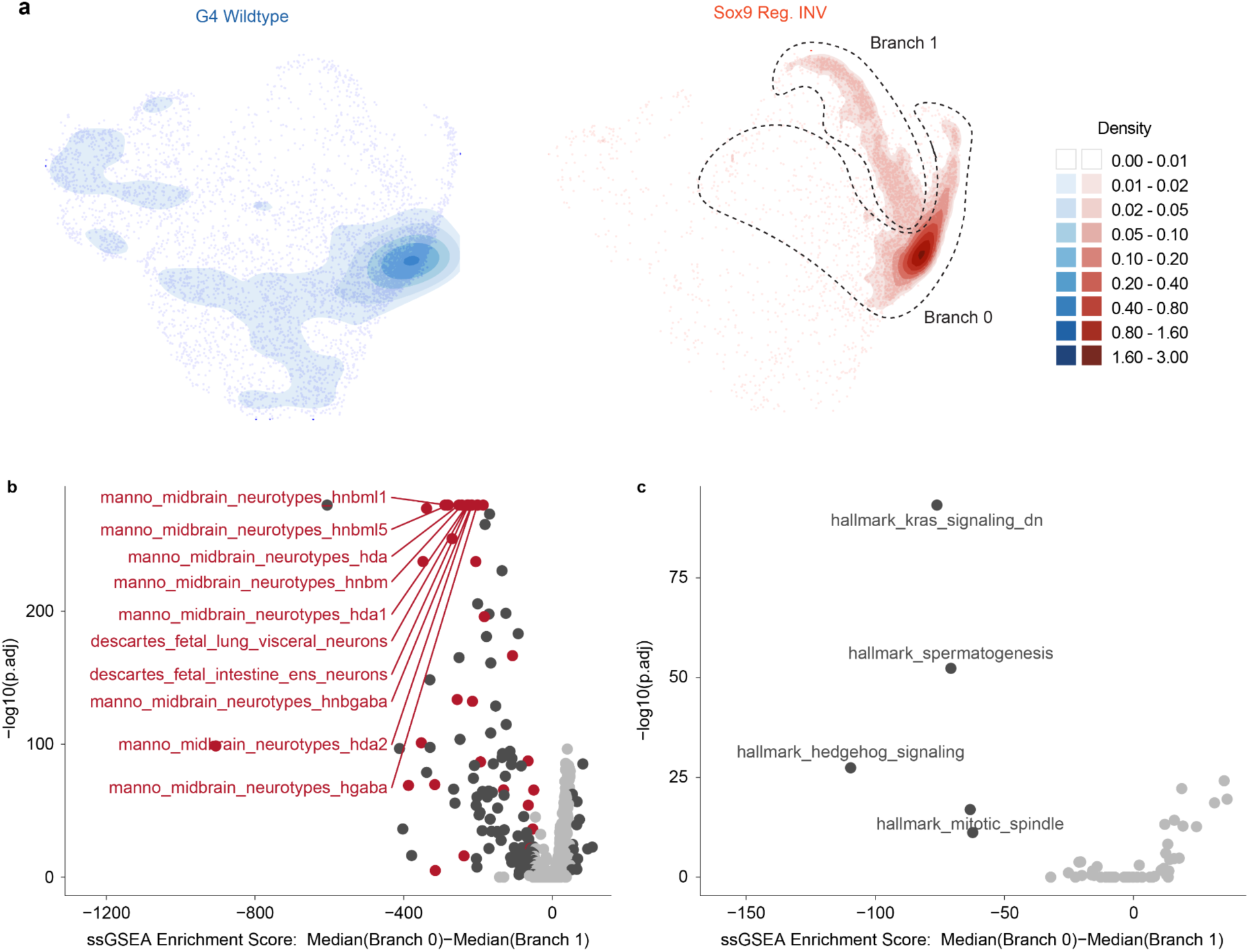
Stalling of Sox9 regulatory INV cells in the undifferentiated mesenchyme and gene set enrichment analysis on these cells. **a**, Density plots for UMAP embedding of G4 wildtype and *Sox9* regulatory INV cells in the limb mesenchymal trajectory (same embedding as **Fig. 4e**). Dotted black lines demarcate the two branches of the undifferentiated mesenchyme, based on the sub-clustering shown in **Fig. 4f. b**,**c,** Comparison of the ssGSEA^84^ scores between the two branches of undifferentiated mesenchyme for *Sox9* regulatory INV cells for (**a**) cell type signature (C8) and (**b**) Hallmark gene sets. Gene sets that are both significantly different between the two branches and that have a difference in median ssGSEA scores greater than 50 are highlighted in dark grey, and the ten most significantly different gene sets are also labelled. In panel **b**, all significantly different gene sets with names containing “neuro” are highlighted in red.

**Supplementary Figure 14.**
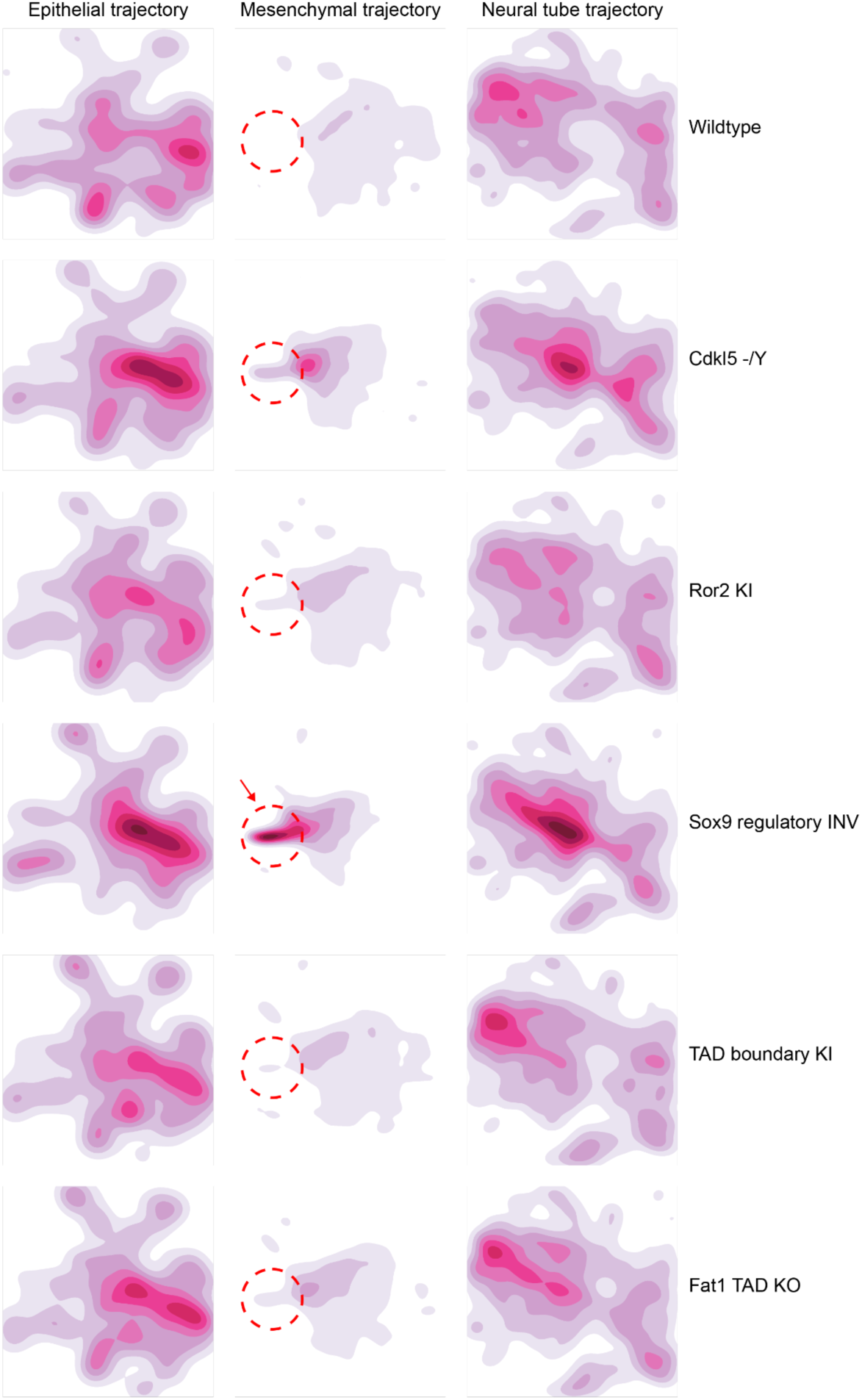
Density plots of the UMAP co-embedding of wildtype and mutant samples from G4 mouse background. We focus on the epithelial, mesenchymal and neural tube main trajectories, which are the three largest. The densities are corrected for the total number of cells. The colour scale is kept consistent across mutants (rows), but varied across the trajectories (columns). Arrow points to the accumulation of cells in the *Sox9* regulatory INV mutant. Dotted circles demarcate the location of cellular accumulation in *Sox9* regulatory INV mutant in the same embedding across all the other mutants.

**Supplementary Figure 15.**
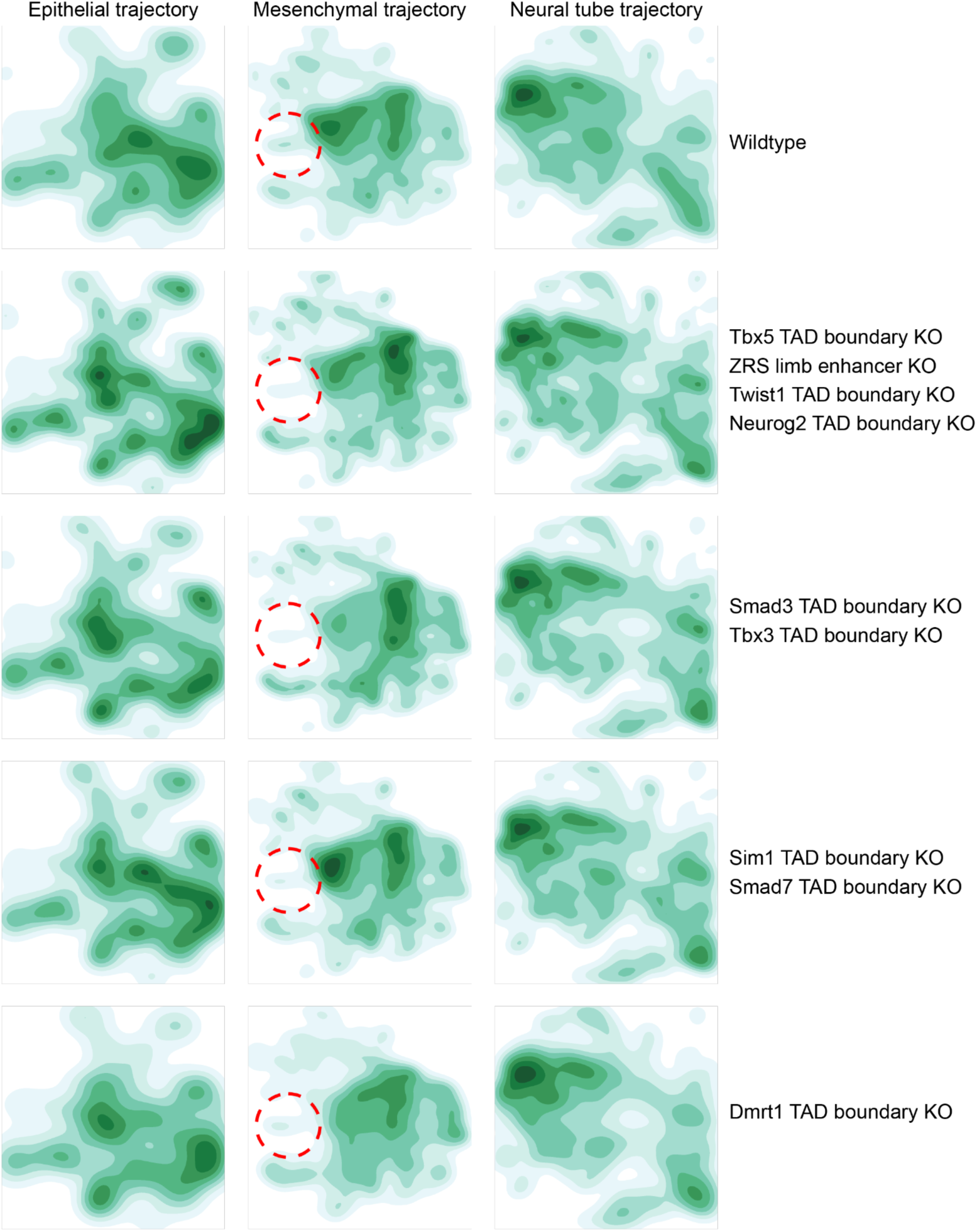
Density plots of the UMAP co-embedding of wildtype and mutant samples from FVB mouse background. We focus on the epithelial, mesenchymal and neural tube main trajectories, which are the three largest. The same embedding as in Supplementary Fig. 14 was used. Mutants with visually similar UMAP embeddings were combined for presentation. The densities are corrected for the total number of cells. The colour scale is kept consistent across mutants (rows), but varied across the trajectories (columns). Dotted circles demarcate the location of cellular accumulation in *Sox9* regulatory INV mutant in the same embedding across all the other mutants.

**Supplementary Figure 16.**
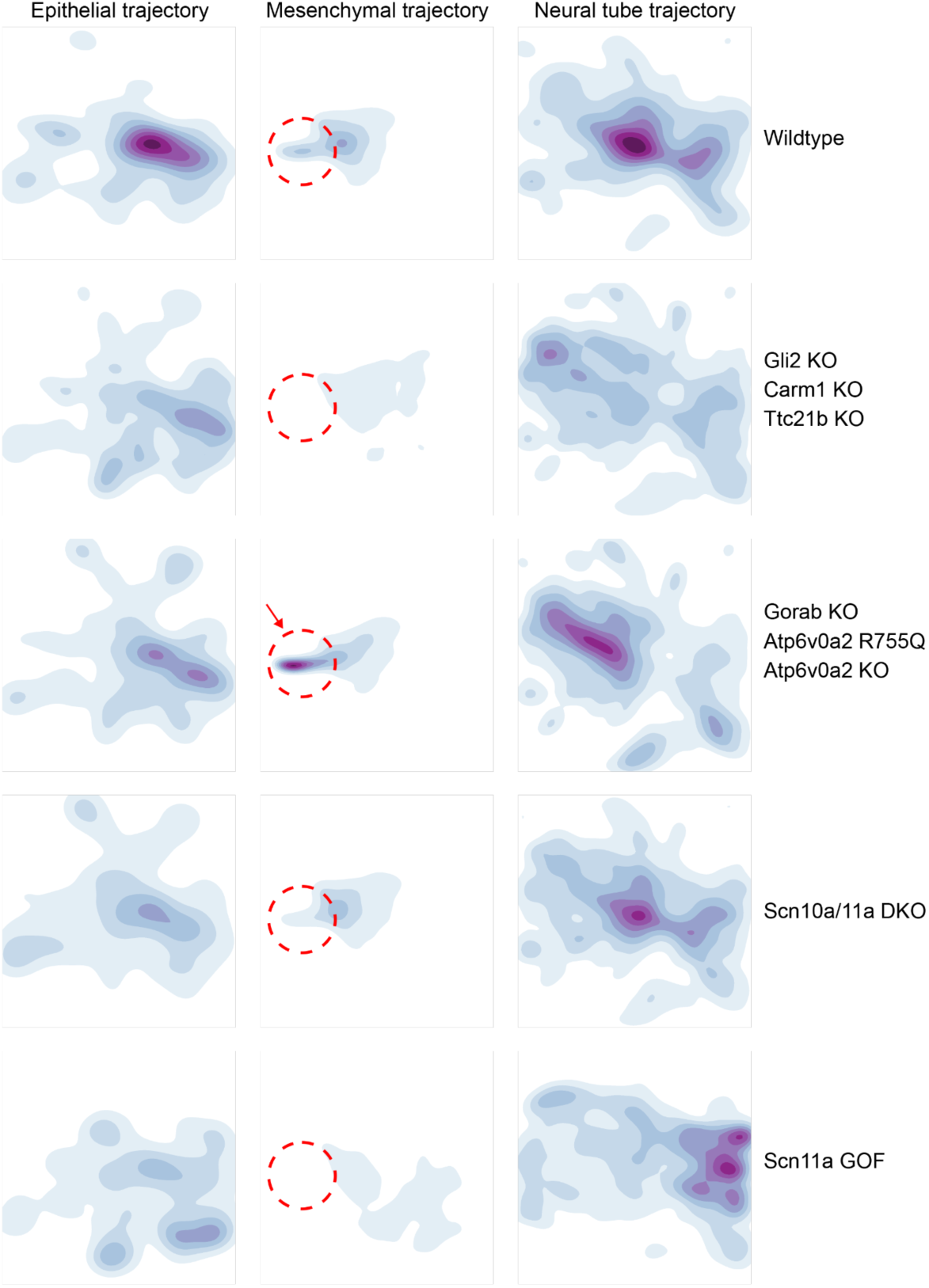
Density plots of the UMAP co-embedding of wildtype and mutant samples from C57BL/6 mouse background. We focus on the epithelial, mesenchymal and neural tube main trajectories, which are the three largest. The same embedding as in Supplementary Fig. 14 was used. Mutants with visually similar UMAP embeddings were combined for presentation. The densities are corrected for the total number of cells. The colour scale is kept consistent across mutants (rows), but varied across the trajectories (columns). Dotted circles demarcate the location of cellular accumulation in *Sox9* regulatory INV mutant in the same embedding across all the other mutants. Arrow highlights a similar accumulation of cells in the *Gorab* KO, *Atp6v0a2* R755Q, and *Atp6v0a2KO* mutants.

**Supplementary Figure 17.**
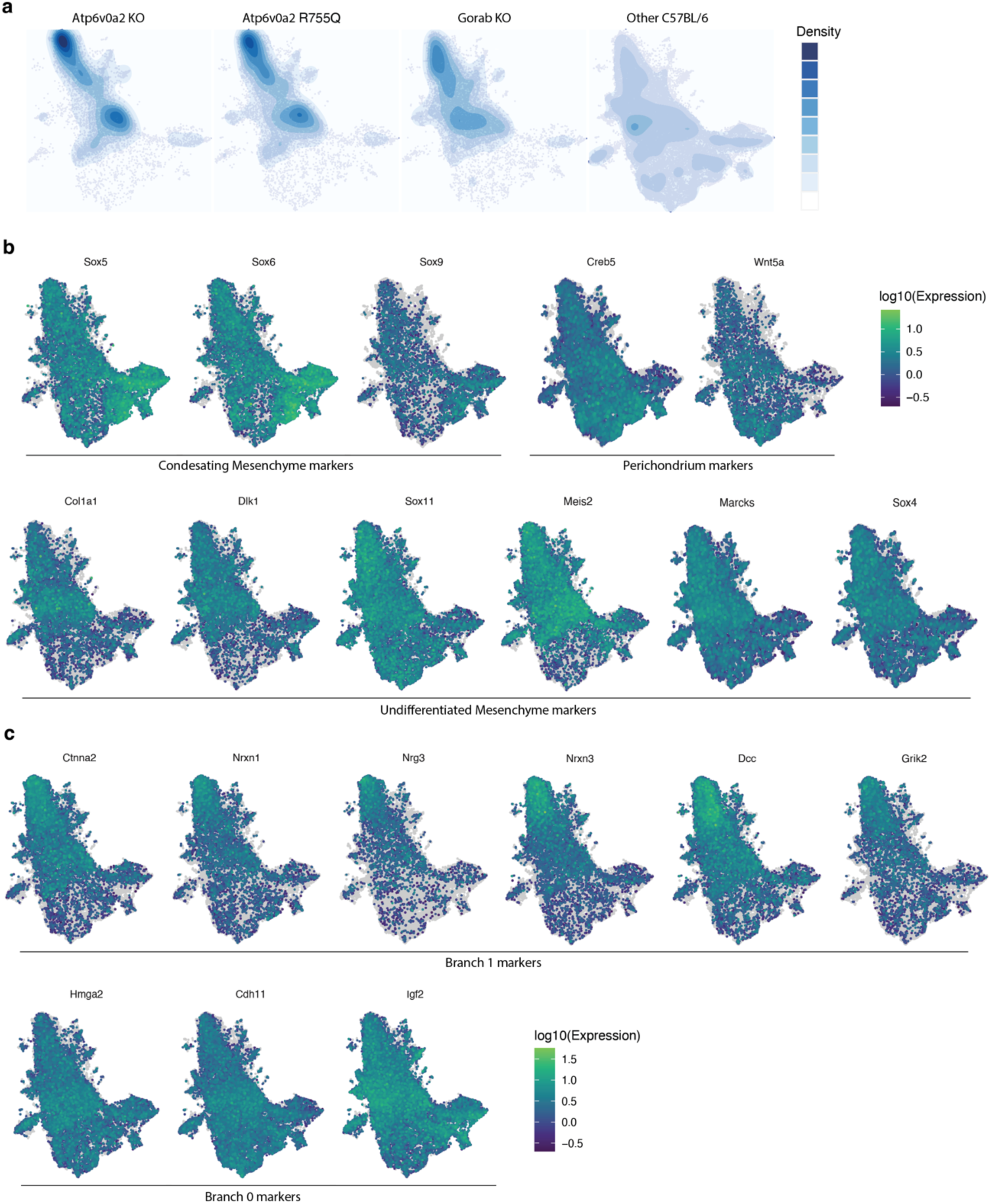
Density and marker gene expression plots of UMAP co-embeddings of wildtype and mutant samples from C57BL/6 mouse background in the limb mesenchyme trajectory. **a,** UMAPs showing the co-embeddings of the limb mesenchyme trajectory for wildtype and mutant genotypes from the C57BL/6 background strain, with cell density and distributions overlaid. **b,** same as in panel **a**, but colored by expression of limb mesenchyme sub-cluster marker genes. The accumulation of cells in the *Gorab* KO, *Atp6v0a2* R755Q, and *Atp6v0a2KO* mutants express markers of undifferentiated mesenchyme. **c,** same as in panel a, but colored by expression of significantly differentially expressed genes between the two branches of Sox9 regulatory INV undifferentiated mesenchyme cells as shown in Fig. 4g.

